# An Open and Continuously Updated Fern Tree of Life

**DOI:** 10.1101/2022.03.31.486640

**Authors:** Joel H. Nitta, Eric Schuettpelz, Santiago Ramírez-Barahona, Wataru Iwasaki

**Author notes:** **Correspondence:** Joel H. Nitta.

## Abstract

Ferns, with about 12,000 species, are the second most diverse lineage of vascular plants after angiosperms. They have been the subject of numerous molecular phylogenetic studies, resulting in the publication of trees for every major clade and DNA sequences from nearly half of all species. Global fern phylogenies have been published periodically, but as molecular systematics research continues at a rapid pace, these become quickly outdated.

Here, we develop a mostly automated, reproducible, open pipeline to generate a continuously updated fern tree of life (FTOL) from DNA sequence data available in GenBank. Our tailored sampling strategy combines whole plastomes (few taxa, many loci) with commonly sequenced plastid regions (many taxa, few loci) to obtain a global, species-level fern phylogeny with high resolution along the backbone and maximal sampling across the tips. We use a curated reference taxonomy to resolve synonyms in general compliance with the community-driven Pteridophyte Phylogeny Group I classification.

The current FTOL includes 5,582 species, an increase of *ca*. 40% relative to the most recently published global fern phylogeny. Using an updated and expanded list of 51 fern fossil constraints, we find estimated ages for most families and deeper clades to be considerably older than earlier studies.

FTOL and its accompanying datasets, including the fossil list and taxonomic database, will be updated on a regular basis and are available via a web portal (https://fernphy.github.io) and R packages, enabling immediate access to the most up-to-date, comprehensively sampled fern phylogeny. FTOL will be useful for anyone studying this important group of plants over a wide range of taxonomic scales, from smaller clades to the entire tree. We anticipate FTOL will be particularly relevant for macroecological studies at regional to global scales and will inform future taxonomic systems with the most recent hypothesis of fern phylogeny.

## 2 Introduction

Ferns (*ca*. 12,000 species) are the second most diverse lineage of vascular plants after angiosperms (*ca*. 300,000 species) and are a useful study system for understanding processes of biogeography (*e.g.*, Tryon, 1986; Kato, 1993), community ecology (*e.g*., Hennequin et al., 2014; Lehtonen et al., 2015), and speciation (*e.g*., Kao et al., 2020). Key to any investigation of evolutionary history in this group is a well-sampled phylogeny. Fortunately, ferns have received relatively intense focus from molecular systematists, which has resulted in the publication of trees for all major clades and DNA sequence data from nearly half of all currently recognized fern species. Thus, there is both a pressing need and sufficient sampling for a globally sampled fern phylogeny.

Past efforts to construct such a global phylogeny have steadily expanded their sampling, at first by mostly generating new sequences, then later by mining GenBank (Pryer et al., 2004; Schuettpelz and Pryer, 2007; Lehtonen, 2011; Testo and Sundue, 2016). Indeed, the growth of plastid fern accessions in GenBank shows no sign of slowing since the most recent global fern phylogeny (Testo and Sundue, 2016; Figure 1). There is a need therefore, not only for a revised global fern phylogeny, but also one that is continuously updated to keep pace with the rapid accumulation of molecular data going forward. Such an effort would eliminate the need for researchers to “reinvent the wheel” each time the need for a globally sampled fern phylogeny arises.

**Figure 1.**
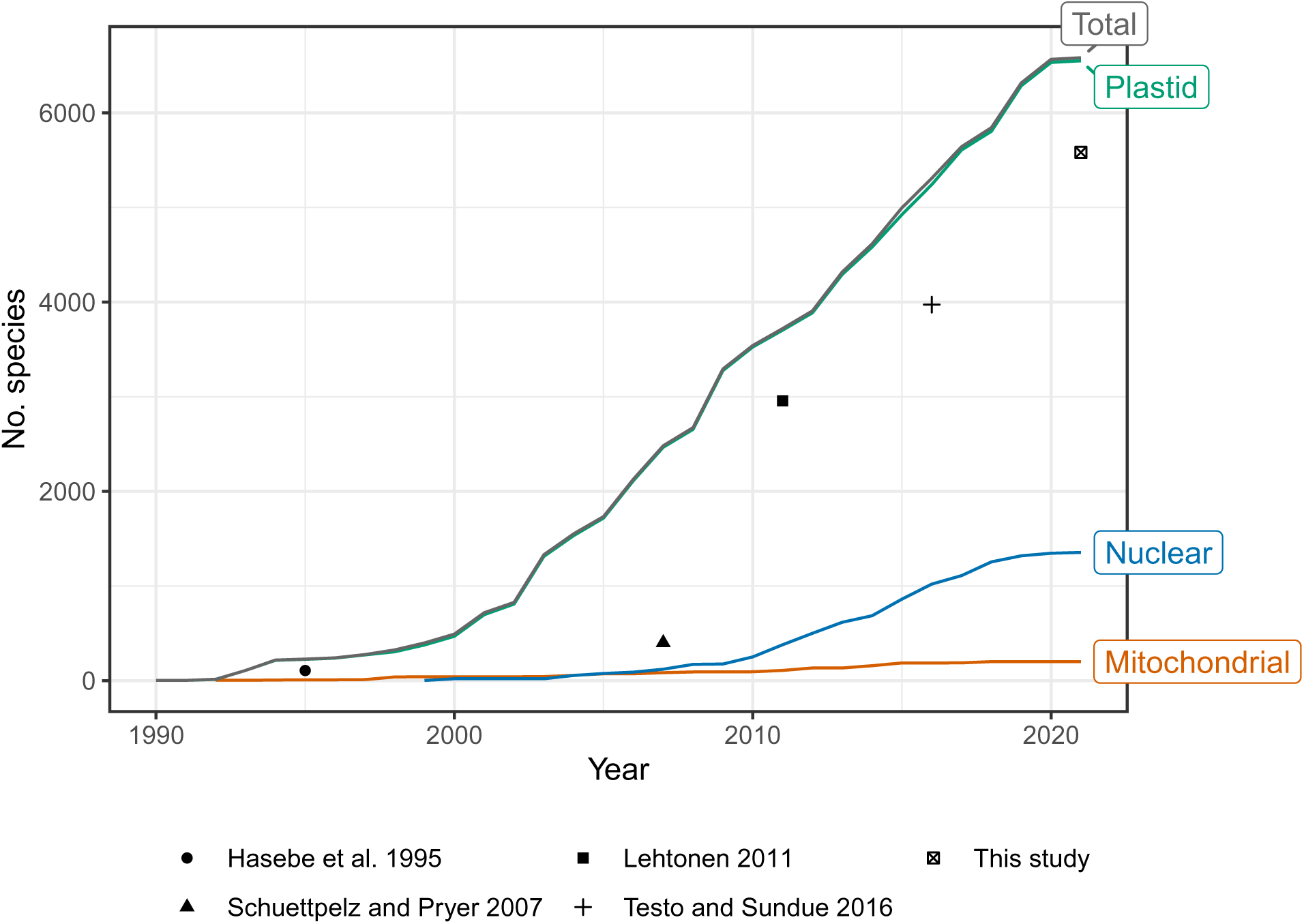
Number of fern species in GenBank by year and genomic compartment, 1990–2021. Points indicate number of species sampled in selected studies of global fern phylogeny (Hasebe et al., 1995; Schuettpelz and Pryer, 2007; Lehtonen, 2011; Testo and Sundue, 2016; this study). This study plotted as 2021 but includes data until 2022-04-15. Schuettpelz and Pryer (2007) did not attempt exhaustive sampling but rather proportional sampling according to lineage size. The relatively small increase in number of species in 2021 may be due to accessions that were still embargoed at the time of writing. Taxonomy of GenBank species follows NCBI (Federhen, 2012). Only accessions identified to species included; environmental samples, hybrid formulas, and names with “aff.” or “cf.” annotations excluded.

Multiple frameworks have been put forth to automatically or semi-automatically generate trees for any particular part of the tree of life (Antonelli et al., 2016), all plants (Eiserhardt et al., 2018), or even the entire tree of life at once (Hinchliff et al., 2015), which would of course subsume a global fern phylogeny. While such approaches are well-suited to some studies, they cannot be expected to produce an optimal fern phylogeny due to the use of “one-size-fits-all” methods to accommodate such a wide phylogenetic breadth. By focusing methods and datasets specifically on ferns, it should be possible to generate a higher quality end-product (tree) that can then be used “as-is” by biologists studying these organisms. Furthermore, there is much to be gained from integrating a carefully designed global fern phylogeny with the fern systematics community that would not be as easily accomplished with a “tree of all life” or “tree of all plants”.

Recently, a taxonomy of ferns and lycophytes was established at the genus level and higher using an inclusive, community-driven approach (Pteridophyte Phylogeny Group I, 2016; hereafter “PPG I”). PPG I has been widely accepted and used, but there were problematic (non-monophyletic) genera included at the time of publication, and many taxonomic changes have been (and will continue to be) proposed since (*e.g*., Almeida et al., 2017; Shang et al., 2018; Zhang et al., 2020). The next iteration of the PPG classification (*i.e*., PPG II) will ideally be an online, open resource that can be updated as necessary. We envision an open, continuously updated global fern phylogeny that could be directly integrated with PPG II such that taxonomic decisions can be made based on community consensus and the most recently available data.

Here, we leverage taxonomic knowledge to design a custom, fully reproducible, mostly automated pipeline to generate a maximally sampled global fern tree of life (FTOL; Figure 2). We plan to run the pipeline on a regular basis and make the results freely available online through a web portal (https://fernphy.github.io) and R package (FTOL working group, 2022b). This will enable anybody interested in the biology of ferns to have access to the most current hypothesis of fern phylogeny, and an associated time-tree dated using a curated list of fern fossils. We anticipate FTOL will have multiple impacts on the field of fern systematics and evolution: 1) it will always provide the most up-to-date snapshot of our collective understanding of fern relationships; 2) it will allow for continuous assessment of taxonomy, and indicate those parts of the tree that are in need of taxonomic revision; and 3) it will be an important source of data for phylogenetic comparative and macroevolutionary studies of ferns.

**Figure 2.**
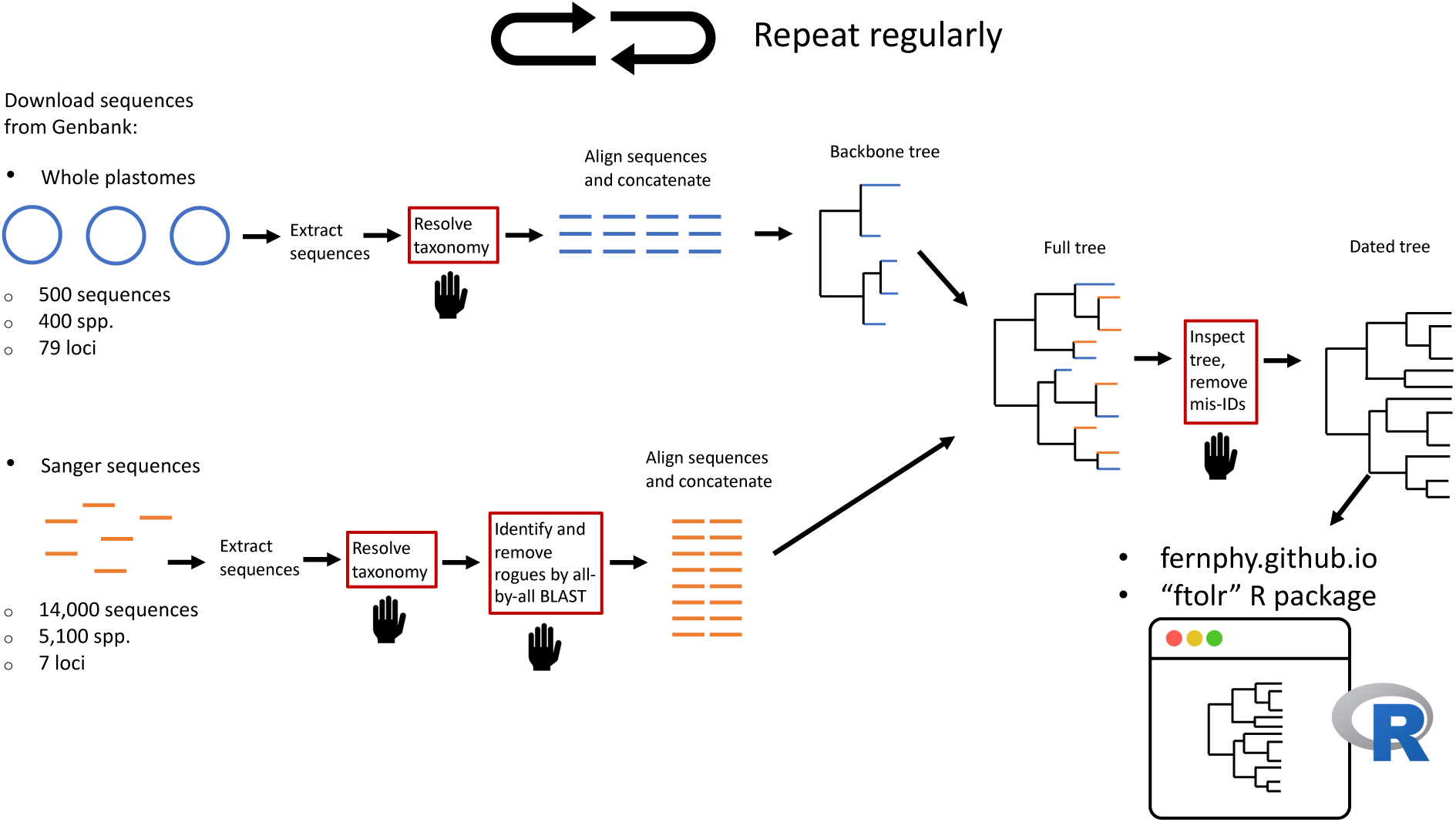
Summary of workflow to construct the Fern Tree of Life (FTOL). The workflow is automated except for steps in boxes with red outlines and a hand symbol. Numbers of sequences and species are approximate. Sequences originating from whole plastomes are in blue; sequences typically obtained by Sanger sequencing are in orange. For details of each step, see Materials and Methods.

## 3 Materials and Methods

### 3.1 Locus selection

The plastid genome has been the most widely sequenced genomic compartment in ferns by far (Figure 1), so we used plastid loci to build our tree. Our sampling includes two major categories of sequence data: 1) seven loci (*atpA, atpB, matK, rbcL, rps4, trnL–trnF*, and *rps4–trnS*) that have been frequently used in molecular analyses of ferns and are typically obtained by PCR and Sanger sequencing (“Sanger loci”); and 2) a much larger set of single-copy loci typically obtained from next-generation sequencing of the plastome (“plastome loci”). Here, the *trnL–trnF* locus includes the *trnL* intron, the 3’ *trnL* exon, and the *trnL–trnF* intergenic spacer; the *rps4–trnS* locus includes only the *rps4–trnS* intergenic spacer. The set of plastome loci is based on the list of 83 protein-coding genes of Wei et al. (2017), which was filtered to only genes that show no evidence of duplication (77 genes, including *atpA*, *atpB*, *matK*, *rbcL*, and *rps4*) and then combined with the *trnL–trnF* and *rps4–trnS* loci (79 loci total). Here, “locus” refers to an individual gene, intergenic spacer, or a unit comprised of these in the case of *trnL–trnF*.

### 3.2 Dataset construction

The “plants” division of GenBank release 249 (cutoff date 2022-04-15) was downloaded from the NCBI FTP server (https://ftp.ncbi.nlm.nih.gov/genbank/). A local database was then created from these data including only ferns and outgroup taxa (select seed plants, lycophytes, bryophytes, and algae) using the restez R package v1.1.0 (Bennett et al., 2018a). GenBank accession numbers corresponding to sequences potentially matching each target locus were obtained by querying GenBank with terms including the locus name (*e.g*., “rbcL”), “Polypodiopsida[ORGN]”, and a date cutoff matching the current GenBank release. Environmental DNA samples and accessions with names including the terms “aff.” or “cf.” or hybrid formulas were excluded (no attempt was made to exclude hybrid taxa with standard names, *i.e*., names that are not formulas). Sequences were then obtained from the local database in FASTA format by GenBank accession number. There is no formal definition of genomic vs. Sanger sequences in GenBank, so we used an empirical sequence length cutoff to distinguish between Sanger (≤ 7,000 bp) and plastome (> 7,000 bp) accessions with the “[SLEN]” term (Figure S1).

There is a lack of consensus on sequence annotation in GenBank; the same locus may be annotated using different names, or not annotated at all. To avoid missing sequences due to differences in annotation format, we used the “Reference_Blast_Extract.py” script of superCRUNCH v1.3.1 (Portik and Wiens, 2020) to extract target loci from GenBank FASTA files. Briefly, this involves querying candidate FASTA files from GenBank with BLAST (Altschul et al., 1997) against a reference database of full length, representative sequences (i.e., a “baited search”; Smith et al., 2009; Smith and Walker, 2019). Those portions of the query that have a significant match in the reference are extracted and written to a new filtered FASTA file.

We constructed the superCRUNCH reference databases by first downloading fern sequences from GenBank, then extracting target gene sequences with a custom R script that parses the GenBank flatfile; this only works for properly annotated accessions. We then filtered the sequences to a single representative longest sequence per genus. Next, we aligned the filtered sequences with MAFFT v7.453-1 (Katoh et al., 2002) and removed poorly aligned regions with trimAl v1.4.rev22 (Capella-Gutiérrez et al., 2009). To maximize the size of the reference database for Sanger loci, we then ran “Reference_Blast_Extract.py” using these sequences as references, followed by filtering to the longest sequence per genus and alignment and cleaning as before; this retrieved additional sequences that lacked annotations in the first round. The cleaned alignments were then used as references for superCRUNCH to obtain a maximally sampled set of fern sequences from accessions downloaded from GenBank.

### 3.3 Taxonomic name resolution

This project aimed to generate a phylogeny that is consistent with PPG I, while accounting for taxonomic changes that have been made since its publication. Species names in GenBank, which use the NCBI taxonomy (Federhen, 2012; Schoch et al., 2020), do not necessarily conform to PPG I. Furthermore, the NCBI taxonomy is not curated specifically for ferns and includes many fern synonyms. Therefore, we standardized all species names in the GenBank sequences (NCBI Taxonomy database dump release 2022-05-01, obtained from https://ftp.ncbi.nlm.nih.gov/pub/taxonomy/taxdump_archive/) against a newly generated reference taxonomy. Our reference taxonomy is based on the World Ferns database v12.8 (Hassler, 2022), which conforms to PPG I (with some exceptions explained below) and is available in Darwin Core format (Darwin Core Task Group, 2009) via the Catalog of Life (Bánki et al., 2021).

To resolve species names, we used the taxastand R package v1.0.0 (Nitta, 2022b), which can account for differences in formatting of taxonomic authors (*e.g*., parenthetical basionym author present or absent) and perform fuzzy matching, which is needed to account for spelling errors or variations in author names.

We manually inspected any fuzzily matched or non-matching names. This revealed some species names in GenBank that were missing in the World Ferns database, spelling errors, and some names in the database that needed to be treated differently (*e.g*., changes in synonymy). We thus updated and edited the initial World Ferns database using the dwctaxon R package v1.0.0 (Nitta, 2022a), which is designed to work with taxonomic data in the Darwin Core format. We refer to the resulting taxonomic database as “pteridocat,” and have made it freely available online (https://github.com/fernphy/pteridocat) so that other researchers may standardize taxonomic names in their data to match those of FTOL (FTOL working group, 2022c).

There are differences between pteridocat and PPG I, mostly at the genus level. In the time since PPG I was introduced, several new genera have been published. Furthermore, multiple genera included in PPG I were known to be non-monophyletic and provisionally circumscribed (Pteridophyte Phylogeny Group I, 2016). pteridocat includes newly published genera, some genera that were not recognized by PPG I, and nothogenera, which were not included in PPG I (Liu et al., 2020). Differences between pteridocat and PPG I are summarized in Table S1.

### 3.4 Removal of rogue sequences using BLAST

Another potential issue with GenBank sequences is the presence of misidentified sequences, poor quality sequences, and contaminants (hereafter collectively referred to as “rogues”). We removed putative rogues from Sanger sequences as follows. First, we constructed a BLAST library including all extracted Sanger sequences using BLAST Suite v2.9.0+. Next, we conducted one BLAST search for each Sanger sequence against the library (all-by-all BLAST). We compared the families (following PPG I) of the matching sequences to each query; in the case the top three best matches belonged to a different family than the query, that query was considered a rogue and excluded from further analysis. For Cyatheales and Saccolomatineae, which include several closely related small families, we used the order and sub-order level, respectively, instead of family to avoid false positives. Species belonging to monotypic families were not considered for this filtering. While this method cannot account for rogues at finer taxonomic levels, it is an efficient approach for removing clearly erroneous sequences. We inspected all accessions flagged this way as rogues before excluding them. In cases where the family mismatch was due to incorrect taxonomy (*e.g*., the species name in GenBank matches the correct family but the name used in the taxonomic database does not), we updated the taxonomic database accordingly.

### 3.5 Sequence concatenation and selection

A typical step in multilocus phylogenetic workflows is to concatenate loci across samples. For phylogenetic studies aiming to include one tip per species, this is often done by concatenating the longest sequence per locus within each species, regardless of source. However, such an approach is potentially problematic because accessions on GenBank may be misidentified and/or species may not be monophyletic. Therefore, we concatenated accessions and selected one final set of concatenated loci per species as follows (here, we refer to “accession” to mean a single locus within a GenBank accession; GenBank accessions may contain multiple loci, but we already split those out using superCRUNCH as described above). First, we constructed gene trees with FastTree v2.1.11-1 (Price et al., 2009, 2010) on default settings, and classified species as monophyletic if they were monophyletic in all gene trees including multiple accessions of that species. Then, we concatenated loci that met any of the following three conditions: 1) if a species was monophyletic, loci were concatenated by selecting the longest accession per locus within that species; 2) if all accessions originated from the same voucher specimen, loci were concatenated across accessions; 3) if all accessions for a given species originated from only one publication, loci were concatenated across accessions. Accessions not meeting any of these conditions were not concatenated. All calculations of sequence length excluded missing bases “?”, or “-”).

We selected the final set of concatenated loci for each species based on presence of *rbcL* and sequence length. We sought to maximize representation of *rbcL* as this is historically the most sequenced locus for ferns, and maximal sampling of one locus has been shown to improve results in super-matrix phylogenies (Talavera et al., 2021). Our procedure for selecting accessions for each species was as follows: 1) if accessions are concatenated including *rbcL* and at least one other locus, select the set of concatenated accessions with the greatest total sequence length; 2) otherwise, if accessions include *rbcL* only, select the accession with the longest *rbcL* sequence; 3) otherwise, select the accession or set of concatenated accessions with the greatest total sequence length. These steps were not needed for plastome data, as each plastome sequence on GenBank originates from a single voucher specimen. For these, we selected the one specimen per species with the longest combined sequence length across all plastome loci.

We also implemented a manual approach for Thelypteridaceae, which had the highest number of non-monophyletic genera (16) in a previous version of FTOL (v1.0.0) and likely has high rates of misidentification in GenBank. This approach used a curated list of accessions from a previous phylogenetic analysis of this family (Patel et al., 2019). After resolving the names in Patel et al. (2019) to pteridocat and correcting some errors (incorrect GenBank accession number in a small number of cases), accessions in this list were used preferentially over those identified by the automated approach.

### 3.6 Sequence alignment

For non-spacer regions (genes), we aligned each locus separately in MAFFT with automatic adjustment for sequence direction and other settings on default. Spacer regions are difficult to align across higher taxonomic levels within ferns (*e.g*., family or higher) as they include frequent indels; however, spacer regions are very useful for phylogenetic analysis at finer scales (*e.g*., within genus or family) where slower-evolving genes like *rbcL* may not provide enough resolution. Therefore, we first aligned spacer regions for each family with MAFFT separately, then merged these subalignments using the MAFFT “--merge” option. This option retains the indels within each subalignment while aligning across subalignments. Sequences from families with fewer than three species each could not be used for subalignments, so these were added as “singletons” during the MAFFT “--merge”. Anemiaceae showed an extremely high number of indels compared to other families and could not be reliably aligned, so we excluded it from the spacer region data (*trnL–trnF* and *rps4–trnS*). We also excluded outgroups from the spacer region data as they cannot be reliably aligned to ferns.

We removed poorly aligned regions, including those with >1% or >5% of sequences having gaps, from the alignments using trimAl (Capella-Gutiérrez et al., 2009).

### 3.7 Phylogenetic analysis

#### 3.7.1 Backbone phylogeny

We generated a backbone phylogeny using maximum likelihood (ML) analysis of the concatenated plastome dataset in IQ-TREE v2.1.3 (Nguyen et al., 2015). ModelFinder (Kalyaanamoorthy et al., 2017) was implemented in IQ-TREE to select the best-fitting model of sequence evolution. To reduce computational burden, we only tested models based on the General Time Reversible (GTR) model (Tavaré, 1986) and did not partition the dataset. The best-fitting model was selected automatically by IQ-TREE according to Bayesian Information Criterion (BIC) score. Node support was assessed with 1,000 ultrafast rapid bootstrap replicates (Minh et al., 2013; Hoang et al., 2018), which were then used to construct an extended, majority-rule consensus tree (the “backbone phylogeny”).

#### 3.7.2 Initial Sanger phylogeny

We used the backbone phylogeny as a constraint tree to conduct initial phylogenetic analysis of the concatenated Sanger dataset in IQ-TREE with the “-fast” option and the GTR+I+G model. We prioritized computational speed for this step, as we needed to repeat it multiple times as described in the next section.

#### 3.7.3 Removal of rogue sequences based on initial Sanger phylogeny

We inspected the initial Sanger phylogeny to identify any remaining rogues or names that needed updating in the taxonomic database. We checked for monophyly at taxonomic levels at or above genus using the MonoPhy R package v1.3 (Schwery and O’Meara, 2016). We updated our taxonomic database if the molecular data clearly indicated the current usage of a synonym was incorrect (*e.g*., a taxonomic intruder into an otherwise expected monophyletic genus with a synonym available for that genus; expected monophyly follows PPG I, 2016). However, this was not possible in all cases, particularly in groups that are in need of taxonomic revision and are known to include non-monophyletic genera (*e.g*., cheilanthoid ferns, grammitid ferns, microsoroid ferns). While we consider our phylogeny may serve as a guide for future taxonomic revisions, we did not make any taxonomic changes that were not already validly published according to the International Code of Nomenclature for Algae, Fungi, and Plants (Turland et al., 2018). Some unpublished names appearing in GenBank and World Ferns databases were not excluded.

Based on the results of this inspection, we updated the list of rogue accessions to be excluded, updated the taxonomic database, and then re-ran all analyses up to this step. During each iteration of the analysis, it is possible that the GenBank accessions added in place of the excluded rogues themselves include rogues. Therefore, we repeated this process until the monophyly of all higher-level (*e.g*., genus rank and higher) taxa that were expected to be monophyletic according to PPG I (2016) was either confirmed or could not be achieved due to outstanding taxonomic issues.

#### 3.7.4 Final Sanger phylogeny

We generated the final Sanger phylogeny using the backbone phylogeny as a constraint tree in ML analysis of the final concatenated Sanger dataset, with the same model selection procedure and bootstrapping as used for the backbone phylogeny. IQ-TREE was initially run using 1,000 iterations (default), but at the end of this run the bootstrap correlation coefficient of split occurrence frequencies was 0.96, which is below the threshold for convergence (0.99). We then continued the search for another 1,000 iterations (2,000 total), but the correlation coefficient fluctuated between 0.96 and 0.98 without overall improvement or convergence. This suggests that the search was stuck in a local optimum. As our alignment contains many species with very similar sequences, it is likely that the search algorithm is unable to optimize many highly similar topologies that only vary in positions of closely related species at the tips. Considering that additional iterations were unlikely to converge, we therefore conducted 10 independent runs of IQ-TREE with 1,000 iterations each, and selected the run with the best (highest) combined log-likelihood of the ML and consensus trees (Zhou et al., 2018).

We did not conduct a more exhaustive analysis testing various partitions of the data or other phylogenetic inference methods or models as our goal is to produce a single species-level phylogeny that is a reasonable hypothesis of fern evolution, not to interrogate the outcomes of multiple, more or less equally applicable methods.

### 3.8 Molecular dating

Dating was conducted separately on the ML tree and the consensus tree using penalized likelihood as implemented in the development version of treePL (commit starting with 551cbde1; Smith and O’Meara, 2012). We initially rooted the tree using *Zygnema circumcarinatum* Czurda, a member of the group of algae (Zygnematophyceae) thought to be most closely related to land plants (Donoghue et al., 2021). Since there is no way to objectively divide branch length between the branch leading to the outgroup and the rest of the tree, we then trimmed *Z. circumcarinatum* from the phylogeny; this effectively positioned bryophytes as the outgroup on a branch or branches with length(s) estimated from the data (Sauquet, 2013).

We selected 51 fern fossils to use as constraints from ferncal v1.0.1, a newly curated database including 145 fern fossil taxa (FTOL working group, 2022a; Table S2, Appendix S2). In the case of redundant fossils (those assignable to the same node in the phylogeny; *e.g*., stems of families that are sister to each other, such as Dipteridaceae and Matoniaceae) we selected the fossil with the oldest age (*i.e*., the upper limit of the oldest stratigraphic period assigned to the fossil). We assigned fossils to lineages only after consulting the original publication (as opposed to simply relying on the fossil name) so we could reassess the identification relative to changes in taxonomy and hypotheses of phylogenetic relationships. Taxonomic concepts may vary between the original publication and currently applied taxonomy (*e.g*., Dicksoniaceae *sensu lato* including other tree fern families used in description of the fossil but Dicksoniaceae *sensu stricto* used currently) and hypotheses of phylogenetic relationships among extant species may change (*e.g*., *Dennstaedtia* used in description of a fossil but this genus now known to be polyphyletic); in both cases, the resulting node to be constrained with a given fossil could change. We applied one fossil constraint outside of ferns (stem euphyllophytes; 407.6 Ma), and fixed the root age of the tree (land plants) at 475 Ma as in Testo and Sundue (2016) and Qi et al. (2018). All constraints other than the root are minimum ages (Figure S2).

We tested rate smoothing parameters in treePL ranging from 1e-12 to 1e-06 (each varies by one order of magnitude); these values were used because our tree spans a wide phylogenetic range with high rate heterogeneity, and because results of initial analyses with higher smoothing values sometimes produced spurious date estimates. A value of 1e-12 was ultimately selected, based on the smallest chi-squared value, for the final analysis.

### 3.9 Reproducibility

The workflow is managed in R v4.2.0 (R Core Team, 2022) with the targets R package v0.12.0 (Landau, 2021). Input data are available at https://doi.org/10.6084/m9.figshare.19474316 (Nitta et al., 2022b). Code used to generate FTOL and compile this manuscript are available at https://github.com/fernphy/ftol and https://github.com/fernphy/ftol_ms, respectively. Docker images to run the analysis and compile this manuscript are available at https://hub.docker.com/r/joelnitta/ftol and https://hub.docker.com/r/joelnitta/ftol_ms, respectively.

## 4 Results

### 4.1 GenBank mining

The initial GenBank query for the seven Sanger loci resulted in 51,226 accessions (note that in some cases a single GenBank accession may contain multiple loci). Extraction of target loci with superCRUNCH recovered 48,719 accessions. Manual inspection of the initial tree resulted in a list of 51 rogues to be excluded. After excluding these and accessions with names that could not be resolved to an accepted name in the taxonomic database, 45,395 accessions were retained. After further excluding rogues identified by the all-by-all BLAST search (Table S3), 45,331 accessions were retained. The final selection of Sanger loci (one set of concatenated accessions per species or one accession per species if conditions for concatenation were not met) after excluding species in the plastome dataset included 13,040 accessions (14,581 sequences when counting distinct loci within each accession separately; Table S4) representing 5,161 species. The most frequent method for joining accessions across loci was by voucher (2,816 species; 54.6%); 1,383 species (26.8%) had accessions that could not be joined or only included one of the seven loci (Table S5).

The initial GenBank query for fern plastomes resulted in 573 accessions, of which 544 accessions were retained after excluding rogues and species with names that could not be resolved. Selection of the longest set of concatenated sequences per species yielded 423 accessions representing unique species (Table S4).

### 4.2 Taxon and locus sampling

Taxon sampling for FTOL v1.1.0 includes 5,582/12,237 species (45.6%), 333/350 genera (95.1%), 48/48 families, and 11/11 orders of ferns (all coverage values are relative to the number of accepted species in pteridocat v1.0.0). Coverage varied by major clade from 29.9% (Gleicheniales) to 69.2% (Osmundales) (Figure 3). Taxon sampling for the backbone phylogeny (derived from plastome loci) includes 423 species (3%), 174/350 genera (50%), 42/48 families (88%), and 11/11 orders of ferns.

**Figure 3.**
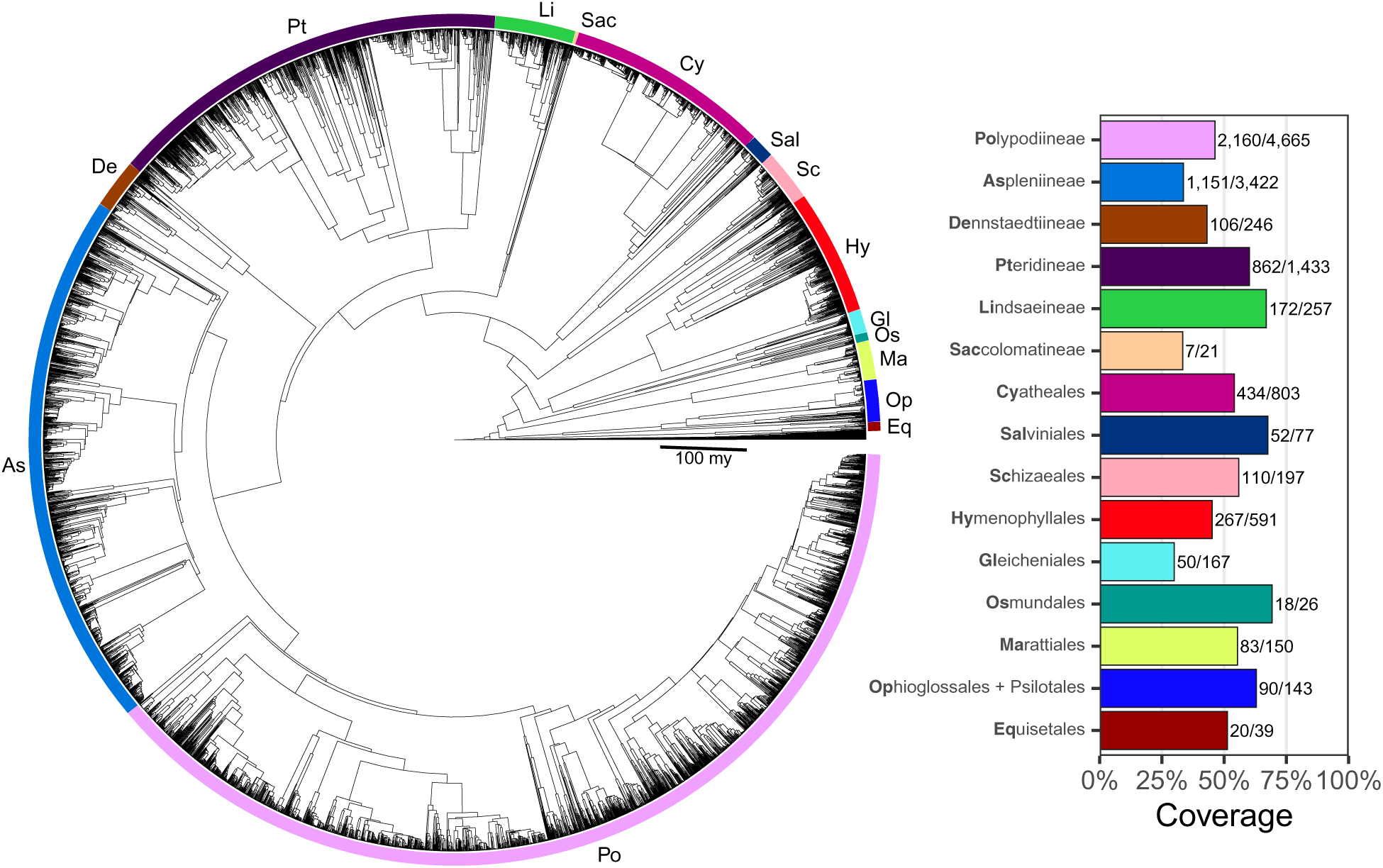
Fern Tree of Life (FTOL). Tree rooted with algae. Inset plot shows coverage by major clade (order or suborder). Bold part of each clade name is its code, which is also indicated on the tree. Numbers next to each bar show sampled species out of total number of species. Taxonomy follows Pteridophyte Phylogeny Group I (2016).

The number of fern species sampled per locus in the Sanger dataset ranged from 1,127 (*matK*) to 4,809 (*rbcL*). A majority of species (4,097; 73.4%) were sampled for more than one locus. A mean of 3.1 ± 1.9 loci were sampled per species (all errors are standard deviations unless otherwise mentioned). The most frequent type of locus sampling per species was *rbcL* alone (1,105 species) (Figure S3). The second most frequent type of locus sampling per species was all seven loci together (558 species). Locus sampling per species in the plastome dataset ranged from 52 to 79 (mean 78.5 ± 2.9 loci per species); 401 species (94.8% of plastome species) included all 79 loci.

### 4.3 DNA alignments

The Sanger DNA alignment was 12,716 bp with 76.9% missing data (missing bases or gaps) overall; rates of missing data by locus ranged from 24.54% (*rbcL*) to 90.18% (*rps4–trnS*). The plastome DNA alignment was 74,883 bp with 12.1% missing data.

### 4.4 Phylogeny

The GTR+F+I+G4 model was selected according to BIC for both the plastome (backbone) and Sanger analyses. Four of the ten runs converged (correlation coefficient of split occurrence frequencies >0.99), but the run with the highest log-likelihood, from which the final tree was selected, did not converge (correlation coefficient 0.976 after 1,000 iterations). The consensus tree had higher log-likelihood (−1,230,734) than the ML tree (−1,230,875), so we only present the topology and divergence times estimated from the consensus tree.

Ferns are strongly supported as monophyletic (BS 100%; all subsequent relationships mentioned received BS ≥ 98% in the backbone phylogeny unless otherwise indicated). The first split within ferns separates a clade including Equisetidae and Ophioglossidae from all other species (Figures 4, S4). The next split separates Marattiales from the remaining ferns, the leptosporangiates (Polypodiidae). Within leptosporangiate ferns, Osmundales is sister to all other species. The next lineage to diverge is a clade including Hymenophyllales and Gleicheniales (BS 98%), followed by Schizaeales. Salviniales was recovered as sister to Cyatheales, which are in turn sister to Polypodiales. The first split within Polypodiales separates a clade including Saccolomatineae and Lindsaeineae from the remainder of species, followed by the subsequent divergences of Pteridineae, then Dennstaedtiineae, which is sister to the eupolypods with moderate support (BS 92%). There are two major clades within eupolypods, Polypodiineae (eupolypods I) and Aspleniineae (eupolypods II).

**Figure 4.**
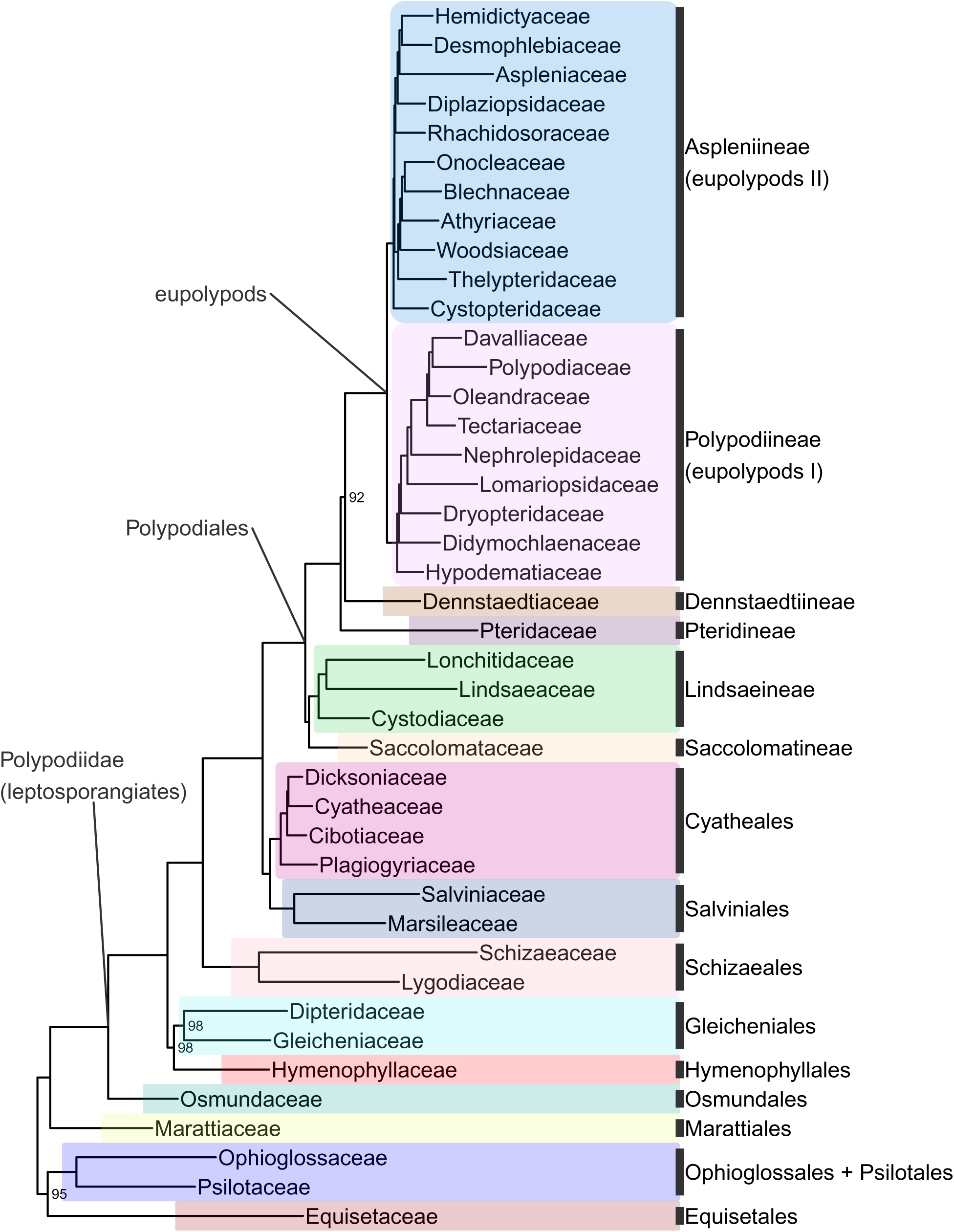
Fern tree of life (FTOL) backbone phylogeny. One exemplar tip is shown per family (all families were found to be monophyletic; see Results). Ultrafast bootstrap support values (%) shown at nodes; unlabeled nodes are 100%. Outgroup (seed plants, lycophytes, bryophytes, and algae) not shown. Colors of major clades (orders or suborders) correspond to those used in Figure 3. Taxonomy follows Pteridophyte Phylogeny Group I (2016); informal clade names in lowercase.

All sampled orders, suborders, families, and subfamilies were recovered as monophyletic (or monotypic) with the exception of Polypodioideae, which is known to be paraphyletic relative to Grammitidoideae (PPG I, 2016). Forty-three genera (13%) were non-monophyletic (Table S6). The subfamilies with the most non-monophyletic genera were Cheilanthoideae (nine), Grammitidoideae (eight), and Thelypteridoideae (seven); other (sub)families each had four or fewer non-monophyletic genera (Table S7).

Bootstrap support was generally moderate to high across the tree (mean 91.7 ±18.1; Sanger phylogeny) and particularly high at deeper nodes (mean 99.7 ± 1.5; 93.2% of nodes with 100% BS; backbone phylogeny). Relationships within some genera were less well-supported, including *Cyathea, Amauropelta*, and parts of *Dryopteris* and *Elaphoglossum* (Figure S5).

### 4.5 Divergence times

We estimate the crown age of ferns to be 423.2 million years (Ma) old. Ages of other major crown groups are: leptosporangiates (392.8 Ma), Polypodiales (303.1 Ma), eupolypods I (161.1 Ma), and eupolypods II (163.0 Ma). Our estimates for leptosporangiates and Polypodiales are both *ca*. 30–40 Ma older than the most recent global fern phylogeny (Testo and Sundue, 2016), while other ages are similar (eupolypods I) or slightly younger (eupolypods II). Estimated stem ages of fern families were mostly older than previous studies (Figure 5). We did not compare crown ages of families across studies because differences in crown ages of smaller clades are likely affected by species sampling.

**Figure 5.**
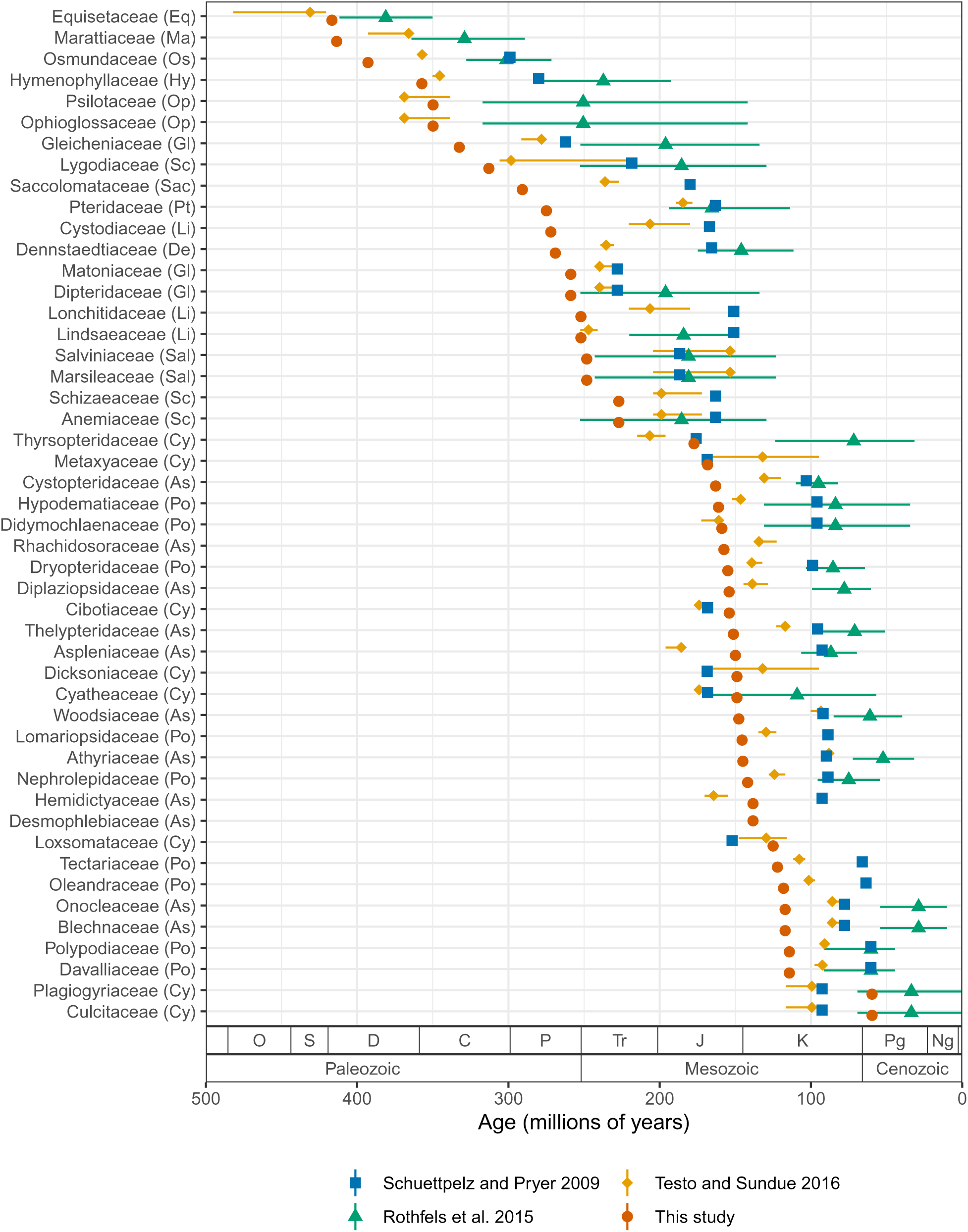
Stem age of fern families (Ma) estimated by selected studies. For studies that used methods with confidence intervals, error bars indicate lower and upper 95% highest posterior density levels and point indicates median (Rothfels et al., 2015; Testo and Sundue, 2016). For other studies, point indicates best (most likely) estimate (Schuettpelz and Pryer, 2009; this study). Codes in parentheses after family names indicate major clade as in Figure 3. Period name abbreviations as follows: O (Ordovician), S (Silurian), D (Devonian), C (Carboniferous), P (Permian), Tr (Triassic), J (Jurassic), K (Cretaceous), Pg (Paleogene), Ng (Neogene).

## 5 Discussion

### 5.1 Nodes of contention in fern phylogeny

The phylogenetic position of Equisetidae relative to other ferns has long been contentious. Here, we recovered Equisetidae as sister to Ophioglossidae (Psilotales + Ophioglossales), in agreement with many other plastid phylogenomic analyses (Grewe et al., 2013; Ruhfel et al., 2014; Gitzendanner et al., 2018; Kuo et al., 2018; Lehtonen and Cárdenas, 2019; One Thousand Plant Transcriptomes Initiative, 2019). However, this contradicts nuclear phylogenomic analyses (Rothfels et al., 2015; Qi et al., 2018; Shen et al., 2018; One Thousand Plant Transcriptomes Initiative, 2019), most plastid analyses with smaller numbers (*ca*. 3–17) of genes (Schneider et al., 2004; Rai and Graham, 2010; Kuo et al., 2011; Testo and Sundue, 2016) and an analysis including mitochondrial data (Knie et al., 2015), which have all recovered Equisetidae as sister to all other ferns. Some earlier plastid analyses based on Sanger data also recovered Equisetidae sister to Marattidae, albeit with generally low support (Pryer et al., 2001; Qiu et al., 2006; Qiu et al., 2007). Aside from alternative resolutions of Equisetidae dependent on genomic compartment or model parameters (Wickett et al., 2014; Kuo et al., 2018), structural support exists for both Equisetidae as sister to Ophioglossidae (*ca*. 550 bp intron in plastid *rps12*; Grewe et al., 2013) and Equisetidae as sister to all other ferns (*ca*. 70 bp intron in mitochondrial *rpl2*; Knie et al., 2015). The clear contradiction between plastid and nuclear phylogenomic data may indicate ancient hybridization or introgression, but robust support for any particular scenario is so far lacking.

Another enigmatic relationship in ferns is the placement of Hymenophyllales. The monophyly of Polypodiidae (leptosporangiate ferns) is well supported across many studies and not in doubt, as is the status of Osmundales as the first lineage to diverge from the remainder of leptosporangiates (Pryer et al., 2001; Schneider et al., 2004; Schuettpelz and Pryer, 2007; Kuo et al., 2011; Testo and Sundue, 2016). However, the subsequent placement of Hymenophyllales differs across studies: some recover Hymenophyllales as the next diverging lineage after Osmundales (Pryer et al., 2001; Schneider et al., 2004; Schuettpelz et al., 2006; Schuettpelz and Pryer, 2007; Testo and Sundue, 2016), while others recover a clade comprising Hymenophyllales sister to Gleicheniales, which then together are sister to the remaining (non-Osmundales) leptosporangiates (Pryer et al., 2004; Lehtonen et al., 2017). Another possibility supported by recent transcriptomic studies is Hymenophyllaceae sister to Gleicheniaceae, which are in turn sister to Dipteridaceae, resulting in a paraphyletic Gleicheniales (Qi et al., 2018; Shen et al., 2018; Shu et al., 2022). Here, we recovered a monophyletic Gleicheniales sister to Hymenophyllales with moderate support, a relationship that was also observed in some, but not all, analyses of plastome data by Lehtonen and Cárdenas (2019) and Kuo et al. (2018). However, our plastome sampling only includes three out of 11 genera of Gleicheniales, and lacks Matoniaceae. While our study was in revision, 42 additional fern plastomes were published, including seven families (*i.e*., Anemiaceae, Culcitaceae, Dipteridaceae, Loxsomataceae, Matoniaceae, Metaxyaceae, and Thyrsopteridaceae) that are not included in our sampling since they were not in the most recent available GenBank release at the time of our analysis (release 249; Du et al., 2022). The addition of Matoniaceae results in non-monophyly of Gleicheniales, with Matoniaceae sister to Dipteridaceae, and these together sister to the remaining leptosporangiates (Du et al., 2022). Furthermore, Du et al. (2022) did not recover a sister relationship between Hymenophyllaceae and Gleicheniaceae. Interestingly, addition of Matoniaceae transcriptome data also supported the non-monophyly of Gleicheniales and the sister status of Matoniaceae + Dipteridaceae in another recent study (Shu et al., 2022).

Within Polypodiales, the relationship between suborders Pteridineae, Dennstaedtiineae and the eupolypods (Polypodiineae and Aspleniineae) has been difficult to resolve (here designated “P”, “D”, and “e”, respectively). Most previous plastid studies based on Sanger sequencing have recovered (D, (P, e)) (Schuettpelz et al., 2006; Schuettpelz and Pryer, 2007; Kuo et al., 2011; Testo and Sundue, 2016; but see Lehtonen, 2011). We recover (P, (D, e)) with moderate support (BS 92%); this topology agrees with other phylogenomic studies based on whole plastomes (Lu et al., 2015; Lehtonen and Cárdenas, 2019) and nuclear data (Rothfels et al., 2015; Qi et al., 2018; Shen et al., 2018), as well as a plastid supermatrix (Lehtonen, 2011). The whole plastome study of Du et al. (2021) recovered a novel topology comprising ((D, P), e) under some analysis settings but (P, (D, e)) under others; a similar study with expanded sampling generally supported ((D, P), e) (Du et al., 2022).

Although some relationships within Polypodiineae (eupolypods I) had previously been resolved differently between various studies using Sanger sequencing, such as Nephrolepidaceae sister to Lomariopsidaceae (Schuettpelz and Pryer, 2007; Zhang and Zhang, 2015) vs. Nephrolepidaceae sister to Tectariaceae, Oleandraceae, Davalliaceae, and Polypodiaceae (Kuo et al., 2011; Lehtonen, 2011; Liu et al., 2013; Testo and Sundue, 2016), our study is in agreement with both nuclear (Qi et al., 2018; Shen et al., 2018) and plastid phylogenomic analyses (Du et al., 2021; Du et al., 2022) that support the latter. Similarly, although Didymochlaenaceae had previously been identified as either sister to the remainder of Polypodiineae (Kuo et al., 2011; Zhang and Zhang, 2015; Testo and Sundue, 2016) or nested with Hypodematiaceae (Schuettpelz and Pryer, 2007; Lehtonen, 2011) by studies using Sanger sequencing, our study as well as nuclear (Qi et al., 2018) and plastid phylogenomic analyses (Du et al., 2021; Du et al., 2022) indicate that Hypodematiaceae is sister to the remainder of the clade.

Relationships of families within Aspleniineae (eupolypods II) have been difficult to resolve due to the ancient, rapid radiation of this clade (Rothfels et al., 2012). Our analysis robustly resolves the relationships between all families in Aspleniineae and is in agreement with a recent plastome analysis with similar sampling (Du et al., 2021). Notably, previous phylogenomic studies that had different or less well-supported topologies did not sample all eupolypod II families (Desmophlebiaceae and Hemidictyaceae absent; Wei et al., 2017; Qi et al., 2018; Shen et al., 2018). The family sister to the remainder of Aspleniineae has been resolved as either Aspleniaceae (*e.g*., Schneider et al., 2004; Testo and Sundue, 2016; Shen et al., 2018) or Cystopteridaceae (*e.g*., Kuo et al., 2011; Wei et al., 2017; Qi et al., 2018). Here, we recovered Cystopteridaceae as sister to the remainder of eupolypod II families, which are split into two clades. One clade consists of Rhachidosoraceae, Diplaziopsidaceae, Aspleniaceae, Desmophlebiaceae, and Hemidictyaceae (RHADD clade of Du et al., 2021; Clade E of Sundue and Rothfels, 2013). The other clade includes Thelypteridaceae, Woodsiaceae, Athyriaceae, Onocleaceae, and Blechnaceae (WOBAT clade of Du et al., 2021; Clade B of Sundue and Rothfels, 2013). Each of these two clades is supported by morphological synapomorphies (Sundue and Rothfels, 2013; Du et al., 2021).

Taken together, our results for nodes of contention in the fern phylogeny generally agree with other plastid phylogenomic analyses and are well within the realm of plausible hypotheses generated to date. We do not consider any of these nodes “solved” by our analysis. Rather, conclusive resolution apparently still awaits additional sampling and perhaps innovation in phylogenetic methods.

### 5.2 Revisiting the timeline of fern diversification

We recovered older crown ages for ferns and large clades therein, and older stem ages for families relative to previous studies (Figures 5, S6). This is almost certainly due to our use of a completely revised and greatly expanded set of fossil calibration points relative to previous studies, which not only resulted in a more densely constrained tree but also a higher number of fern families with minimum fossil ages. Our set of fossil calibration points did not add any extremely old fossils (> 200 Ma) that would be expected to strongly push back ages across the tree (save for stem Marattiaceae, which is well-known for its extensive fossil record; Rothwell et al., 2018); rather, the vast majority of newly added calibration points are younger than 150 Ma (Figure S7).

Several recent studies exploring divergence times across a global fern phylogeny all used a similar set of 24–26 fossil calibration points (Schuettpelz and Pryer, 2009; Testo and Sundue, 2016) or secondary calibration points based on studies using this set (Rothfels et al., 2015). However, there were a few inconsistencies in the application of these fossils due to differences in taxonomic concepts between the original fossil publication and the studies in which they were used. More importantly, our set of calibration points approximately doubles the number used by previous studies and it is likely that this expanded dataset is responsible for the older stem age estimates for many families (Figures 5, S6). To test this hypothesis, we conducted an additional analysis using the fossil constraints of Testo and Sundue (2016) but otherwise the same methods (FTOL analyzed with treePL; Figure S8). The resulting stem family ages show a much closer agreement with those of Testo and Sundue (2016) (*R*^2^ = 0.89, *P* = 5.79e-23, linear model; Figures S9, S10), indicating that our expanded set of fossil calibration points, not differences in topology or dating methodology, is the primarily contributor to the older ages observed in the current study.

Notably, the scenario of fern diversification suggested by our results somewhat conflicts with the hypothesis that Polypodiales diversified “in the shadow of angiosperms” (Schneider et al., 2004; Schuettpelz and Pryer, 2009). Rather, we estimate that the origin (*i.e*., stem age) of many polypod families coincides with or even precedes the diversification and rise to ecological dominance of angiosperms during the Late Cretaceous (Benton et al., 2022) (Figure S11). Our estimated age of (303.1 Ma) for crown Polypodiales is considerably older than other recent studies (Testo and Sundue, 2016; Du et al., 2021) and the fossil record, which only dates back to the Early Cretaceous (Chen et al., 1997; Schneider and Kenrick, 2001; Deng, 2002; Schneider et al., 2016; Regalado et al., 2018). Molecular ages that are significantly older than the fossil record should be treated with caution; yet, our study is in agreement with others in suggesting a “long fuse” between initial appearance of Polypodiales and their subsequent diversification and widespread preservation in the fossil record (Testo and Sundue, 2016; Du et al., 2021).

Due to the large size of our dataset, carrying out more detailed molecular dating analyses (*e.g*., Bayesian analysis) is computationally difficult with currently available methods (*e.g*., BEAST; Drummond and Rambaut, 2007; Bouckaert et al., 2014). Here, we have prioritized computational speed and simplicity, since we anticipate re-running the pipeline on a regular basis. We therefore consider our dated tree as a starting point, and not the final word, for a re-evaluation of divergence times in ferns. Future studies should focus on utilizing our greatly expanded fern fossil dataset to conduct more thorough molecular dating analyses, possibly including alternative schemes for the age of the root and testing the effects of different parameters used for setting priors in Bayesian analyses.

### 5.3 Plastid vs. nuclear fern trees

Plastid sequences are convenient for phylogenetic analysis because they are essentially a single, uniparentally inherited linkage group, thus free from recombination. However, a tree derived from plastid data may not necessarily mirror those inferred from other data sources. Conflict between plastid and nuclear phylogenies has been frequently observed in narrowly focused (*e.g*., genus level) studies using traditional Sanger sequencing (*e.g*., Sessa et al., 2012; Zhou and Zhang, 2017; Wei et al., 2021) and has recently been demonstrated at deeper levels within Polypodiaceae using phylogenomic approaches (Wei and Zhang, 2022). Such conflict does not necessarily reflect insufficient methodology or sampling, but rather may be due to processes including (but not limited to) introgression, lineage sorting, and hybridization at deep phylogenetic levels. Therefore, a major future research goal for fern molecular systematics should be to combine nuclear and plastid datasets to infer species trees with comprehensive sampling.

Recent transcriptomic studies are gradually clarifying the backbone of the fern phylogeny using many (25–2,400) nuclear genes from representative species spanning the tree (Rothfels et al., 2015; Qi et al., 2018; Shen et al., 2018). The most comprehensively sampled phylogenomic study targeting ferns is the on-going Genealogy of Flagellate Plants (GoFlag) project, which seeks to generate genomic data (*ca*. 300 single-to-low copy nuclear gene regions) for all flagellate plants (bryophytes, lycophytes, ferns, and gymnosperms; Breinholt et al., 2021). GoFlag data have recently been used in a phylogenomic analysis and taxonomic revision of Thelypteridaceae resulting in the recognition of multiple new genera (Fawcett et al., 2021; Fawcett and Smith, 2021), and additional phylogenomic analyses of other fern groups using GoFlag markers are to be expected in the near future.

The rapid growth of genomic data notwithstanding, species level sampling of such nuclear phylogenomic datasets is still far less than that available from the plastome (Figure 1). Furthermore, many subclades of ferns are under active investigation using both Sanger and next-gen sequencing of plastid markers, and plastid data for previously unsampled species will likely continue to grow at a rapid pace. We therefore expect that the methodology outlined here will continue to be useful to generate a maximally sampled plastid fern tree of life for many years to come.

### 5.4 Comparison with other automated phylogeny pipelines

Ferns are not unique in having large amounts of publicly available DNA sequence data, and several pipeline tools exist that can leverage such data to automatically generate phylogenies (Pearse and Purvis, 2013; Xu et al., 2015; Antonelli et al., 2016; Bennett et al., 2018b; Drori et al., 2018; Smith and Walker, 2019; Portik and Wiens, 2020). In particular, superSMART (Antonelli et al., 2016) and pyPHLAWD (Smith and Walker, 2019) are recently developed pipelines that can generate maximally sampled phylogenies for any higher taxon of choice. The latter was used to generate a broadly inclusive seed plant phylogeny (Smith and Brown, 2018).

Although we make use of some of the functionality of these tools (*e.g*., restez for generating a local copy of GenBank; superCRUNCH for identifying orthologous sequences; Bennett et al., 2018a; Portik and Wiens, 2020), our pipeline is mostly custom-built for ferns. We chose our approach because it allowed us to implement various steps specific to ferns that would not be possible using a fully automated, taxon-agnostic pipeline like superSMART or pyPHLAWD, which we believe ultimately results in a higher-quality fern phylogeny. For example, we integrate our phylogeny with a custom taxonomy for ferns (pteridocat), tailor alignment strategies to locus and taxon (fern-wide alignment for coding loci, nested alignment within families for non-coding loci), inspect intermediate results and modify input based on expert taxonomic knowledge (curation of sample inclusion and exclusion lists), and employ a custom set of criteria for selecting and concatenating sequences within species. While we have not sought to make our workflow available as a general tool due to its high degree of specialization, some of the methods employed here could be adapted and implemented in other phylogeny pipelines.

### 5.5 Accesibility and usage of FTOL

We have sought to make FTOL easily available to support research on the evolution and ecology of ferns. FTOL is available via the ftolr R package (FTOL working group, 2022b), and a web portal (https://fernphy.github.io). Furthermore, we have made all the underlying data (*e.g*., DNA alignments, fossil calibrations) available so that other researchers can use these to conduct analyses such as further investigations of divergence times or phylogenetic analysis including custom sets of DNA sequences (*e.g*., Nitta et al., 2022a).

A typical step in any analysis that joins data across multiple sources (*e.g*., trait data and a phylogeny) is to resolve taxonomic names so that the usage of synonyms does not prevent data merging (Page, 2008). During the preparation of FTOL, we developed two additional R packages that enable taxonomic name resolution to join data with FTOL: pteridocat (FTOL working group, 2022c) and taxastand (Nitta, 2022b). We selected R because it is widely used by the biological research community, well established, and freely available (Lai et al., 2019). The pteridocat package includes the pteridocat taxonomic database as a data frame (tibble) in Darwin Core format. The taxastand package includes functions to resolve taxonomic names while taking into account variation in taxonomic author format and orthographic variation. By using these two packages in combination, it should be straightforward for other researchers to map their own data onto FTOL, thus greatly enabling and enhancing studies including, but not limited to, comparative phylogenetics, biogeography, and community ecology in ferns.

### 5.6 FTOL as a living, community-driven resource

We want to be clear that FTOL is in no way meant to be the “official” fern phylogeny; it is simply one reasonable hypothesis that has been designed to be maximally inclusive at the species level. FTOL cannot substitute for careful systematic studies at finer taxonomic scales that include sampling of multiple individuals per species and/or other sources (*e.g*., morphological, nuclear) of data, and such studies continue to be vital to our understanding of fern evolution.

One feature of FTOL that sets it apart from the vast majority of other phylogenetic studies is its iterative nature. Unlike most other published fern phylogenies, the current version of FTOL described in this manuscript is not meant to be the last. Rather, we plan to re-run these analyses as additional data become available on GenBank, and release updated versions of the tree on a regular basis. We envision that FTOL will be integrated with the next iteration of the Pteridophyte Phylogeny Group classification, PPG II, to provide the most recent hypotheses on the monophyly of various fern taxa, which will in turn enable a more natural classification system.

Furthermore, FTOL will not only grow in size with time, but also become more refined. We are aware that our methodology cannot produce a “perfect” tree, nor is that our goal. Indeed, we anticipate that there will almost always be tips in the tree that need correction, either because they get overlooked (*e.g*., placement of species within genera, which we did not have the resources to inspect) or because an updated taxonomic treatment is not yet available (*e.g*., cheilanthoid ferns). One example of progressive refinement is Thelypteridaceae, which had the highest number of non-monophyletic genera (16) in the previous version of FTOL (v1.0.0). After implementing a manual inclusion list for Thelypteridaceae and consulting with a taxonomic expert on this family (S. Fawcett), the number of non-monophyletic genera was reduced to seven. While manual inclusion lists are not an ideal long-term solution because they cannot grow with GenBank, this case demonstrates that such lists are a reasonable option for taxonomically difficult groups, and more importantly, the improvement that can result from input from taxonomic experts. Another example of the advantage of a continuously updated tree is the publication of 42 additional fern plastome samples while this paper was in revision (Du et al., 2022). While we were unable to integrate these data into the current version of FTOL, they will be automatically added when the next version of GenBank data is released, and we expect the next version of FTOL will reflect this new knowledge.

To facilitate community-driven improvements, we have made our methodology (code), data, and software (R packages and docker image) completely open and available. It is our hope that other researchers using FTOL will contribute by making edits and suggestions, preferably through the GitHub repository (https://github.com/fernphy/ftol). This way, FTOL will continually improve and keep pace with the currently available data and taxonomic hypotheses of ferns.

## Acknowledgments

The authors thank M. Hassler for maintaining the World Ferns taxonomic database and making it available to use for research. Members of the Iwasaki lab provided comments that improved the manuscript. A.E. White provided helpful comments on an early version of the analysis. S. Fawcett provided helpful comments on taxonomy of Thelypteridaceae.

