## Supplementary figures and images for "An Open and Continuously Updated Fern Tree of Life"

### Figure S2

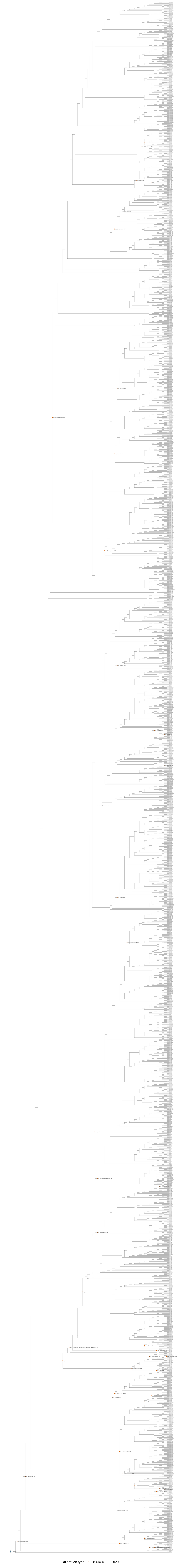

### Figure S4

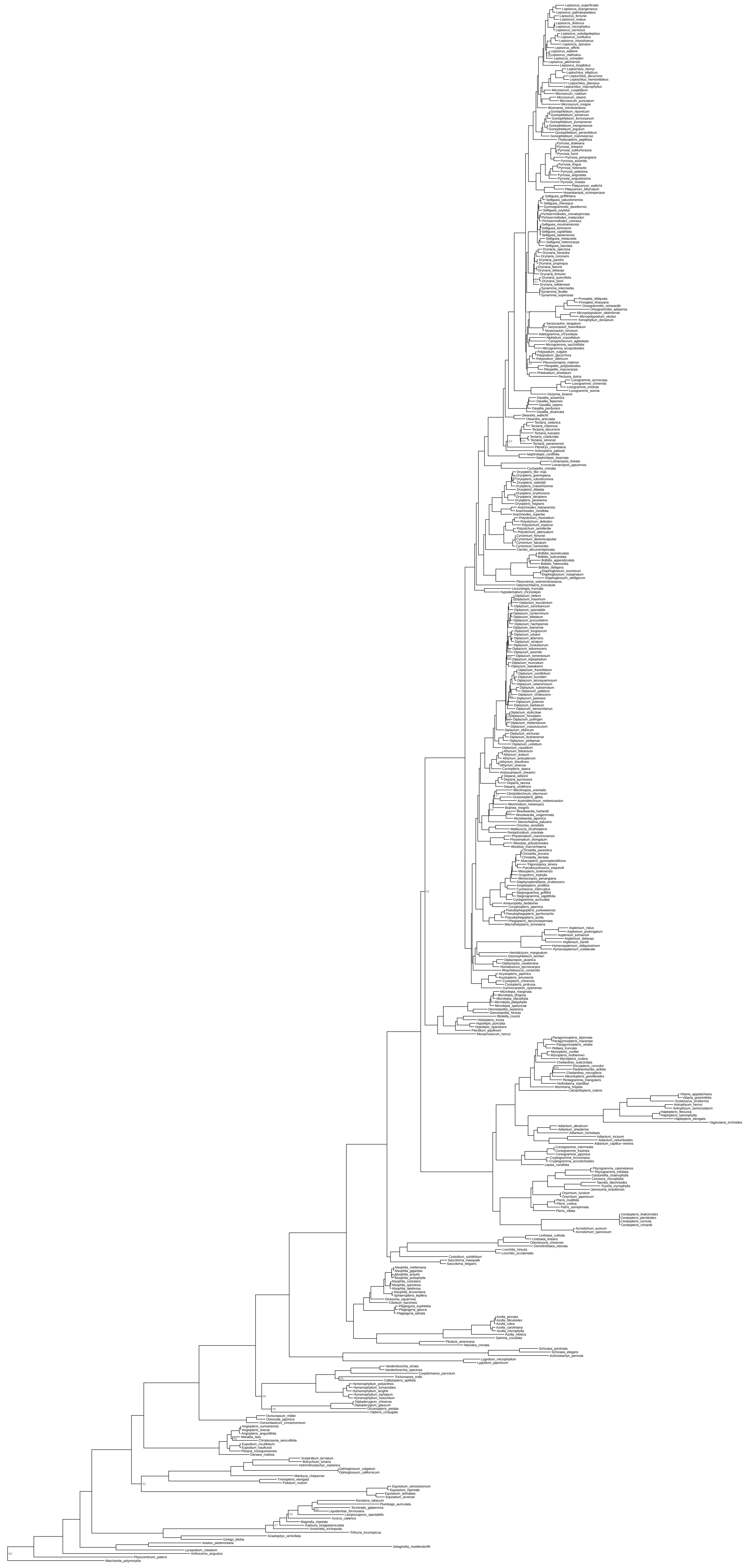

### Figure S5

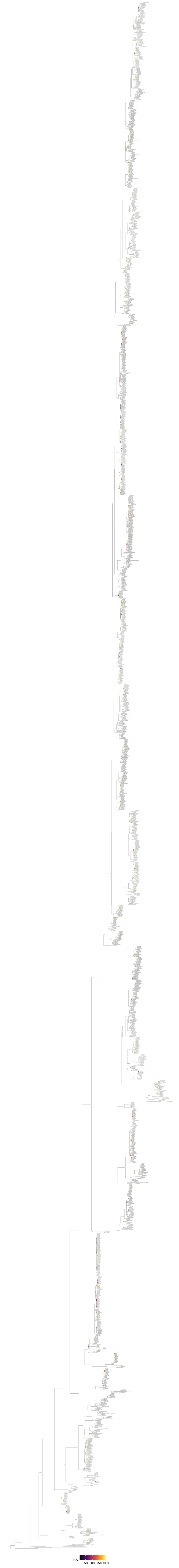

### Figure S8

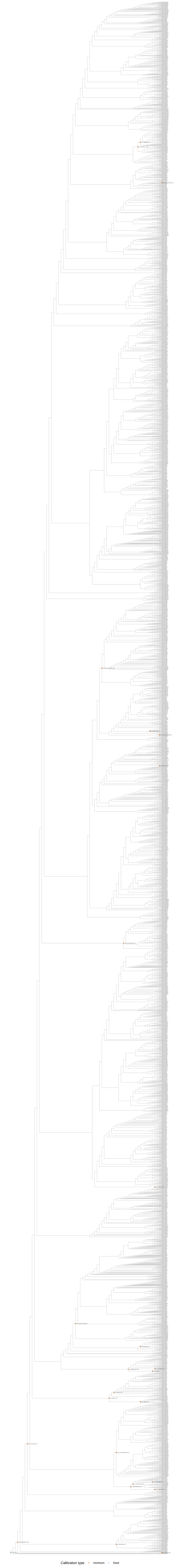
