## Appendix A1 for "An Open and Continuously Updated Fern Tree of Life"

Tables and Figures

**Table S1**. Differences in genera included as accepted in "pteridocat" taxonomic database v1.0.0 (https://github.com/fernphy/pteridocat) vs. PPGI (Pteridophyte Phylogeny Group I, 2016) (Table_S1.csv). "diff" is type of difference (pteridocat genus not in PPGI or PPGI genus not in pteridocat). "n_species" is the number of accepted species in each genus (pteridocat) or estimated number of species (PPGI). References in "note" column provided at end of this appendix. "World Ferns" refers to Hassler (2022), which was modified to create pteridocat.

**Table S2**. Fossils used as calibration points for molecular dating (Table_S2.csv). "n_fos" is the fossil ID number used in the "ferncal" database v1.0.1 (https://github.com/fernphy/ferncal). "minimum_age" is the age used for calibration (Ma); "lower" is the lower limit of the oldest stratigraphic age for the fossil (Ma); "upper" is the upper limit of the oldest stratigraphic age for the fossil (Ma). For detailed notes and full references for each fossil, see Appendix A2.

**Table S3**. GenBank accessions identified as rogues by all-by-all BLAST (Table_S3.csv). "q_family" is the family of the GenBank query (accession). "s_family" is the family of the top three best matches (only cases where the top three matches all belong to the same family included). All accessions in this table were excluded from further analysis.

**Table S4**. GenBank accessions used in this study (Table_S4.csv). "species" is the species name as it appears in the phylogenetic tree. "seq_len" is the total number of bases excluding missing bases ("?", "N", or "-") and only includes bases belonging to loci used in this study (not the entire accession). "sci_name" is the scientific name in the pteridocat taxonomic database. "ncbi_name" is the scientific name used in the NCBI taxonomic database version 2022-05-01. "ncbi_taxid" is the NCBI taxonomy database unique identifier. "outgroup_status" is "FALSE" for ingroup taxa (ferns) and "TRUE" for outgroup taxa.

| Method | n | % |
|---|---|---|
| Publication | 138 | 2.67% |
| Manual | 166 | 3.22% |
| Monophyletic | 658 | 12.75% |
| Unjoined | 1,383 | 26.80% |
| Voucher | 2,816 | 54.56% |
| Total | 5,161 | |

**Table S5**. Methods used to join accessions across loci for Sanger dataset. “manual” only used for Thelypteridaceae (see Methods). Count is number of species.

| Taxonomic level | Monophyly | n | % |
|---|---|---|---|
| Order | Yes | 11 | 100% |
| Suborder | Yes | 6 | 100% |
| Family | Monotypic | 4 | 8.3% |
| Family | Yes | 44 | 91.7% |
| Subfamily | Monotypic | 2 | 8.0% |
| Subfamily | Yes | 22 | 88.0% |
| Subfamily | No | 1 | 4.0% |
| Genus | Monotypic | 67 | 20.1% |
| Genus | Yes | 223 | 67.0% |
| Genus | No | 43 | 12.9% |

**Table S6**. Monophyly by taxonomic level for ferns in FTOL.

**Table S7**. Non-monophyletic fern genera in FTOL (Table_S7.csv). “number_tips” is the number of tips (species) in the genus. “delta_tips” is the number of tips that do not belong to the genus but are included in the clade descending from the most common recent ancestor of the genus. “number_intruders” is the number of intruding taxa into the genus. “number_outliers” is the number of tips (species) belonging to the genus that appear in other clades. Status of intruders and outliers was determined using the default cutoff level (0.5) in the R package “MonoPhy” (Schwery and O’Meara, 2016).

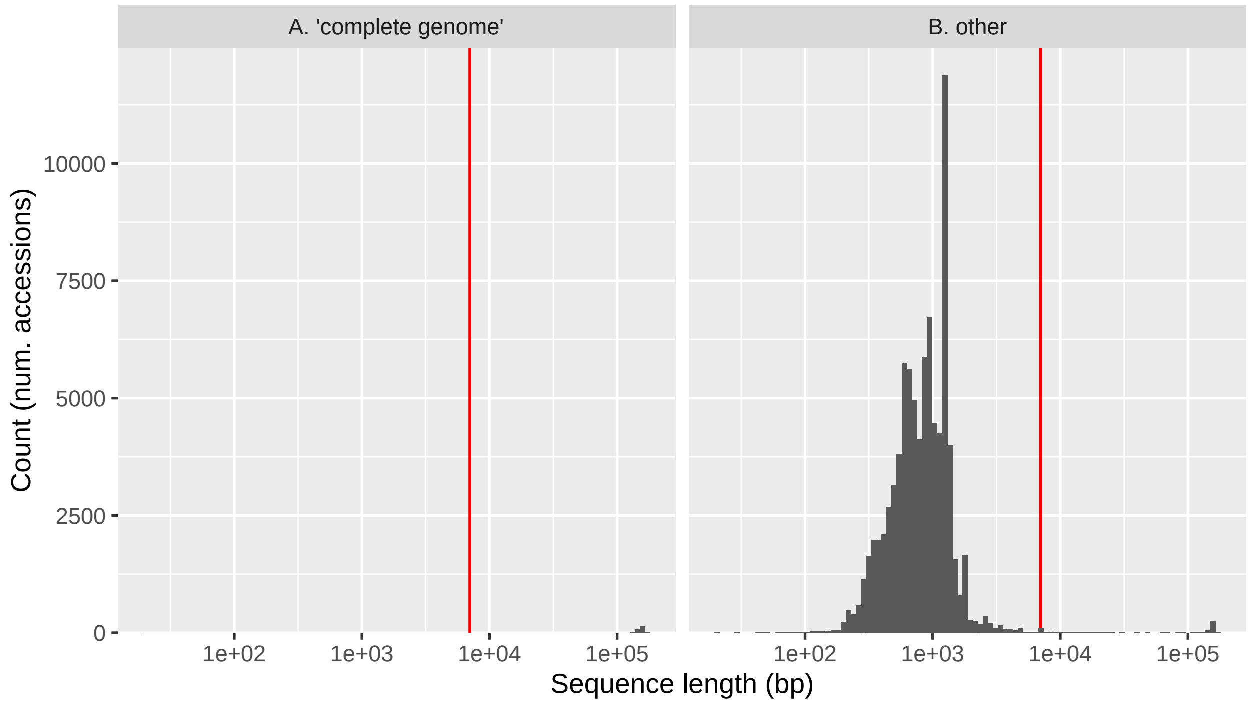


**Figure S1**. Histogram of sequence lengths (bp; log-scale) of fern accessions in GenBank release v249. Fern accessions obtained by querying GenBank for "Polypodiopsida[ORGN]" with length 10–200,000 bp. **A.** Sequences with the term 'complete genome' in their description (*e.g.*, "Angiopteris evecta chloroplast, complete genome"). **B.** Other sequences (those without 'complete genome' in their description). Vertical red line at 7,000 bp indicates empirical length cutoff used to differentiate between between Sanger (≤ 7,000 bp) and plastome (> 7,000 bp) accessions. Note that "other" includes some whole plastome accessions that cannot be distinguished based on description alone (e.g., "UNVERIFIED: Nephrolepis biserrata plastid sequence"); these are mostly above 7,000 bp.

**Figure S2**. Consensus phylogeny (Sanger dataset) with node labels indicating placement of fossil constraints (Figure_S2.pdf). Number for each constraint is age in Ma. Outgroup taxon *Zygnema circumcarinatum* (algae) pruned prior to dating analysis and not shown.

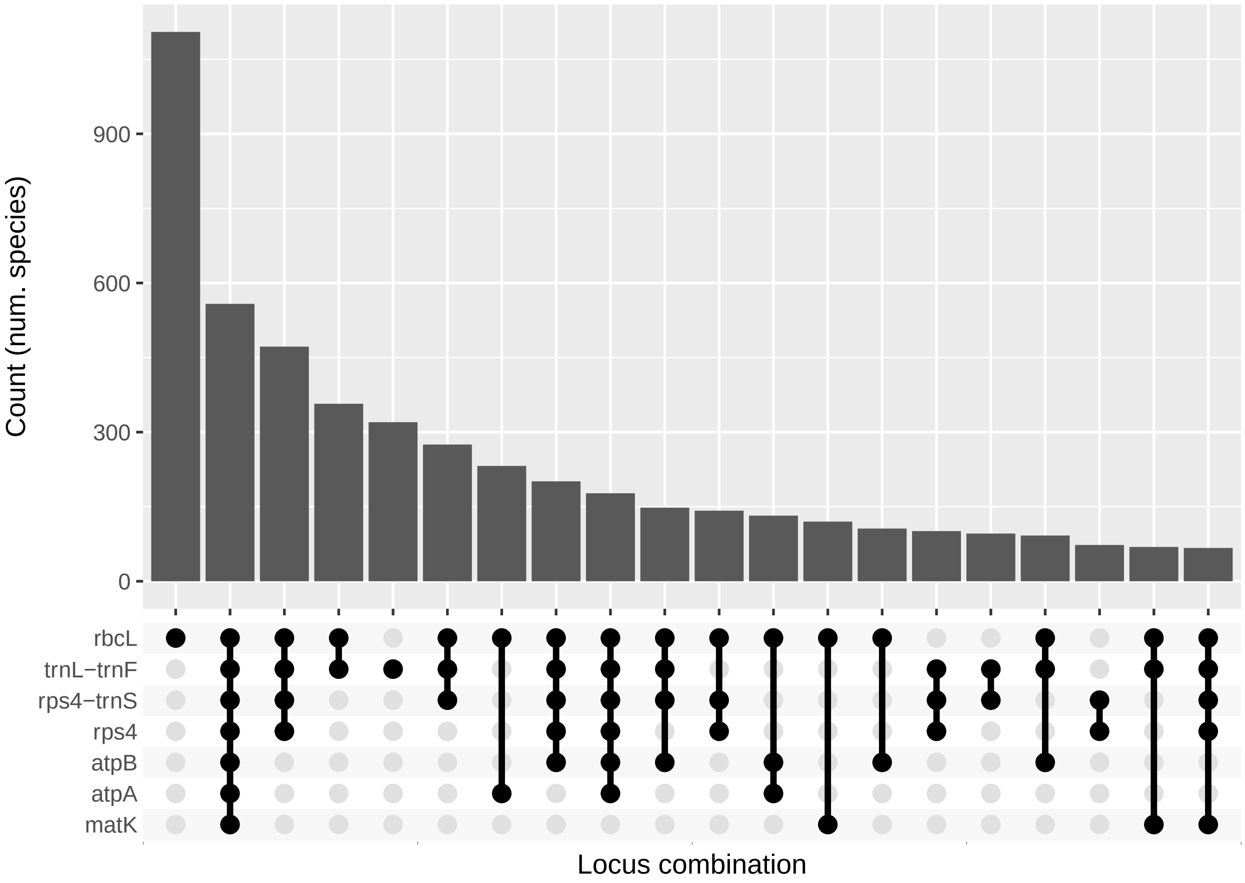


**Figure S3**. Frequency of locus combinations by species, Sanger dataset. Only top 20 combinations shown.

**Figure S4**. Consensus plastome (backbone) phylogeny (Figure_S4.pdf). Values at nodes are bootstrap support (%); only values less than 100% are shown. Outgroup taxon *Zygnema circumcarinatum* (algae) not shown.

**Figure S5**. Consensus phylogeny (Sanger dataset) with node labels showing bootstrap support (%) (Figure_S5.pdf). Darker colors indicate lower support. The backbone (plastome) phylogeny was used as a constraint tree, so all clades present in the backbone phylogeny tree receive 100% bootstrap support. Outgroup taxon *Zygnema circumcarinatum* (algae) not shown.

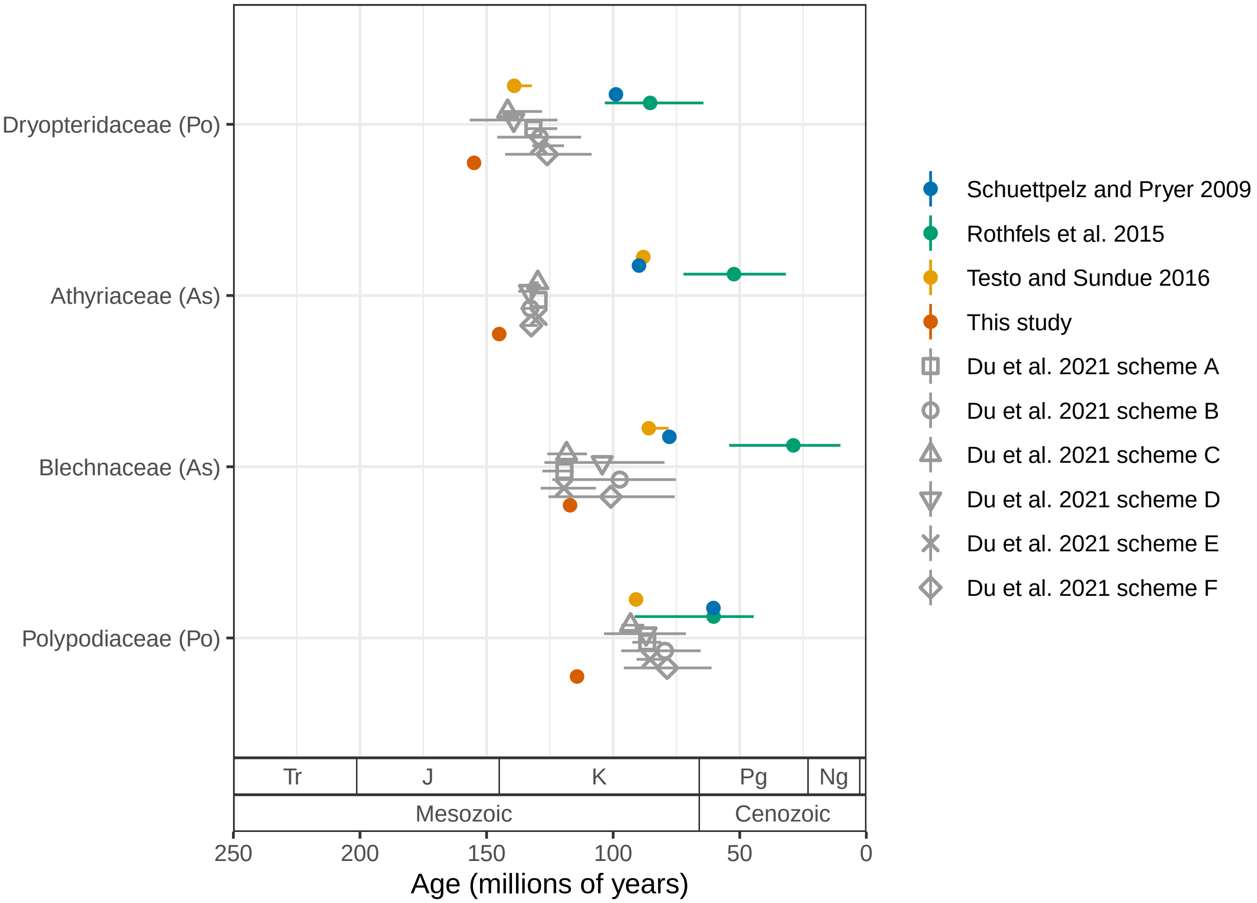


**Figure S6**. Stem age of fern families (Ma) estimated by selected studies, only including families with stem ages that were provided across all studies. For studies that used methods with confidence intervals, error bars indicate lower and upper 95% highest posterior density levels and point indicates median (Rothfels et al., 2015; Testo and Sundue, 2016; Du et al., 2021). For other studies, point indicates best (most likely) estimate (Schuettpelz and Pryer, 2009; this study). Du et al. (2021) used six different dating schemes; for details, see that paper. Codes in parentheses after family names indicate major clade as in Figure 3. Period name abbreviations as in Figure 5.

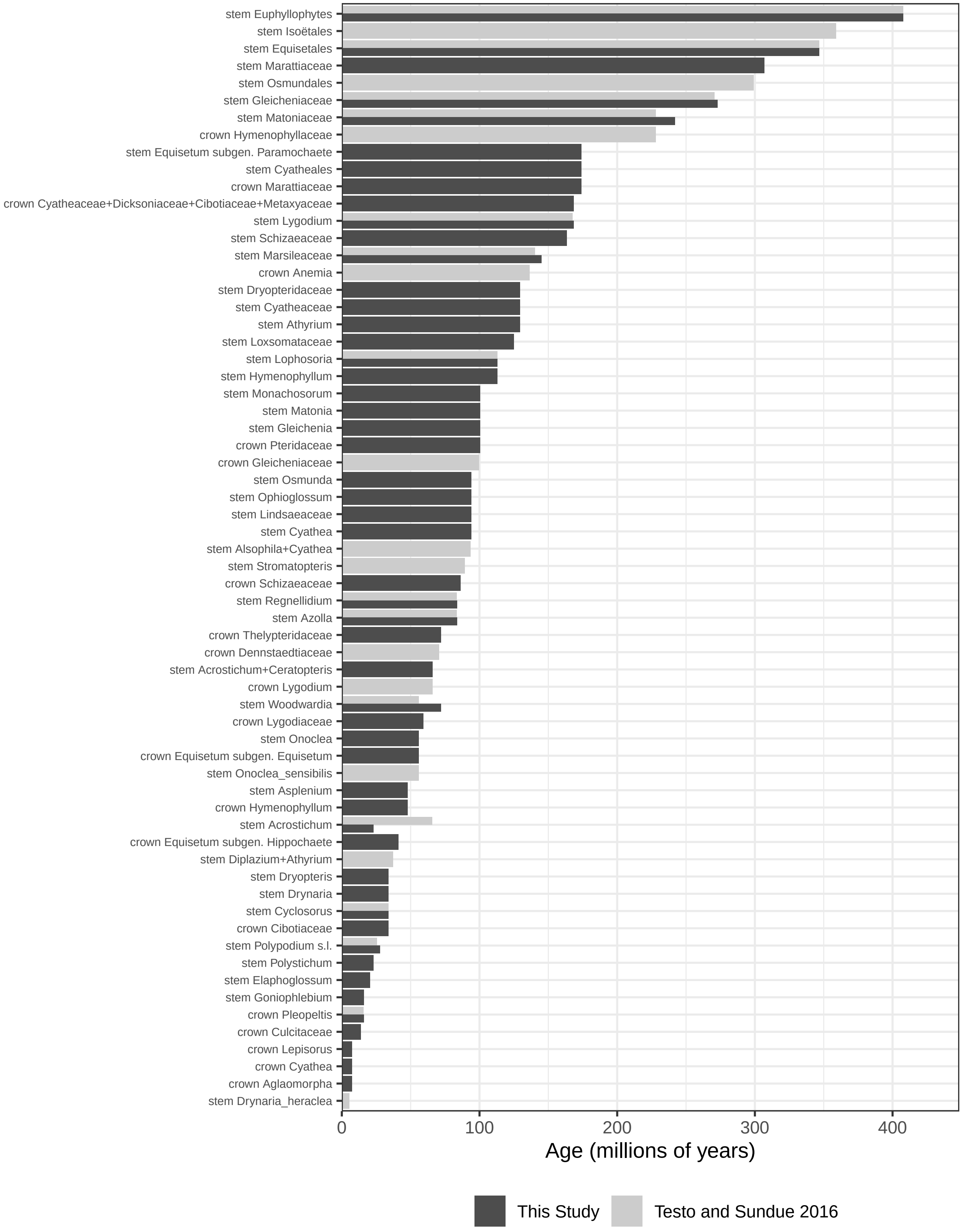


**Figure S7**. Age of nodes constrained by fossils in this study vs. Testo and Sundue (2016). Wide bars indicate the node was only constrained in one study; narrow bars indicate the the node was constrained in both studies.

**Figure S8**. Consensus phylogeny (Sanger dataset) with node labels indicating placement of fossil constraints of Testo and Sundue (2016) (Figure_S8.pdf). Number for each constraint is age in Ma. Outgroup taxon *Zygnema circumcarinatum* (algae) pruned prior to dating analysis and not shown.

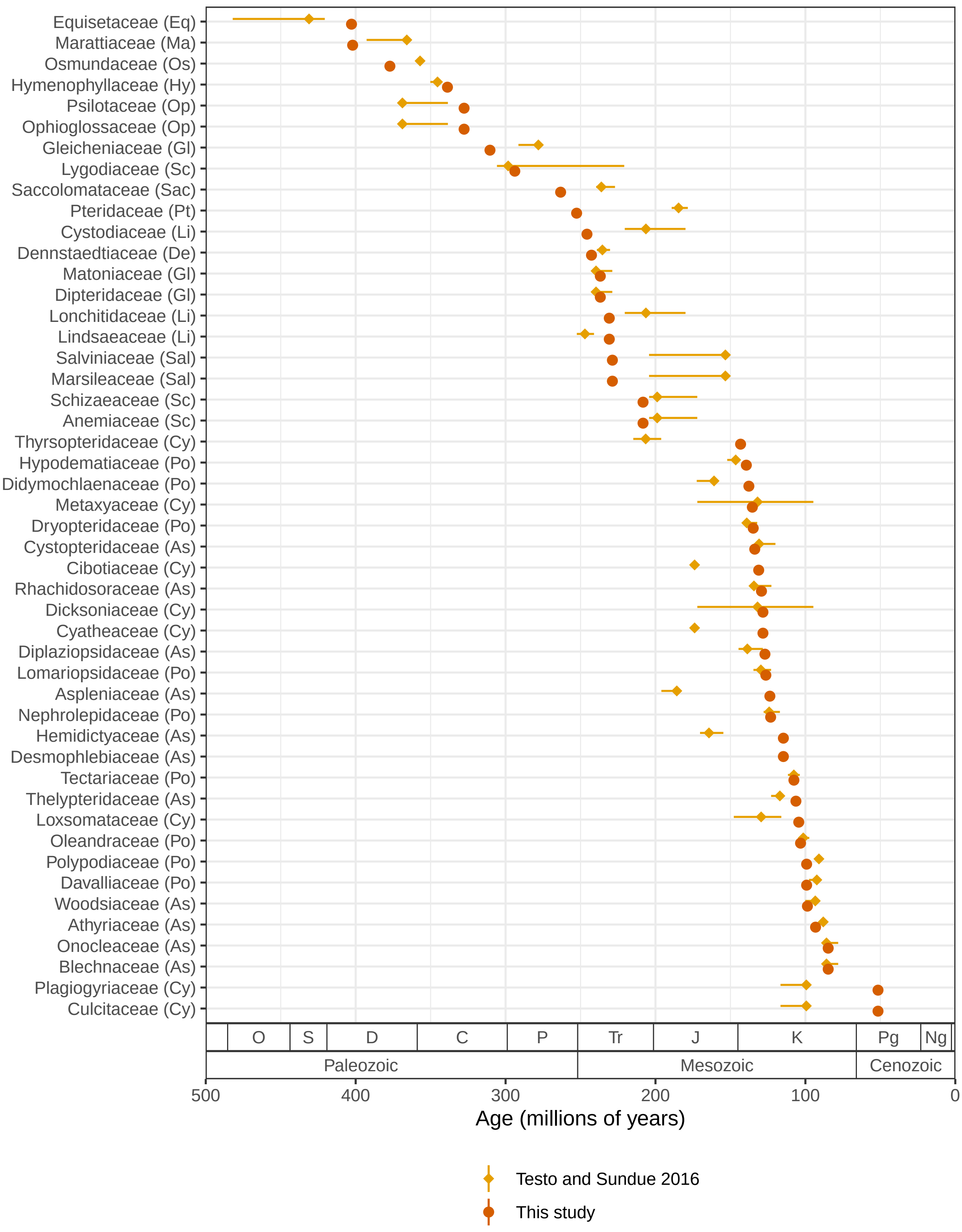


**Figure S9**. Comparison of stem age of fern families (Ma) estimated by Testo and Sundue (2016) and those estimated using methodology (treePL) and phylogeny (FTOL) of the current study with the fossil constraints of Testo and Sundue (2016). For Testo and Sundue (2016), error bars indicate lower and upper 95% highest posterior density levels and point indicates median For this study, point indicates best (most likely) estimate. Codes in parentheses after family names indicate major clade as in Figure 3. Period name abbreviations as in Figure 5.

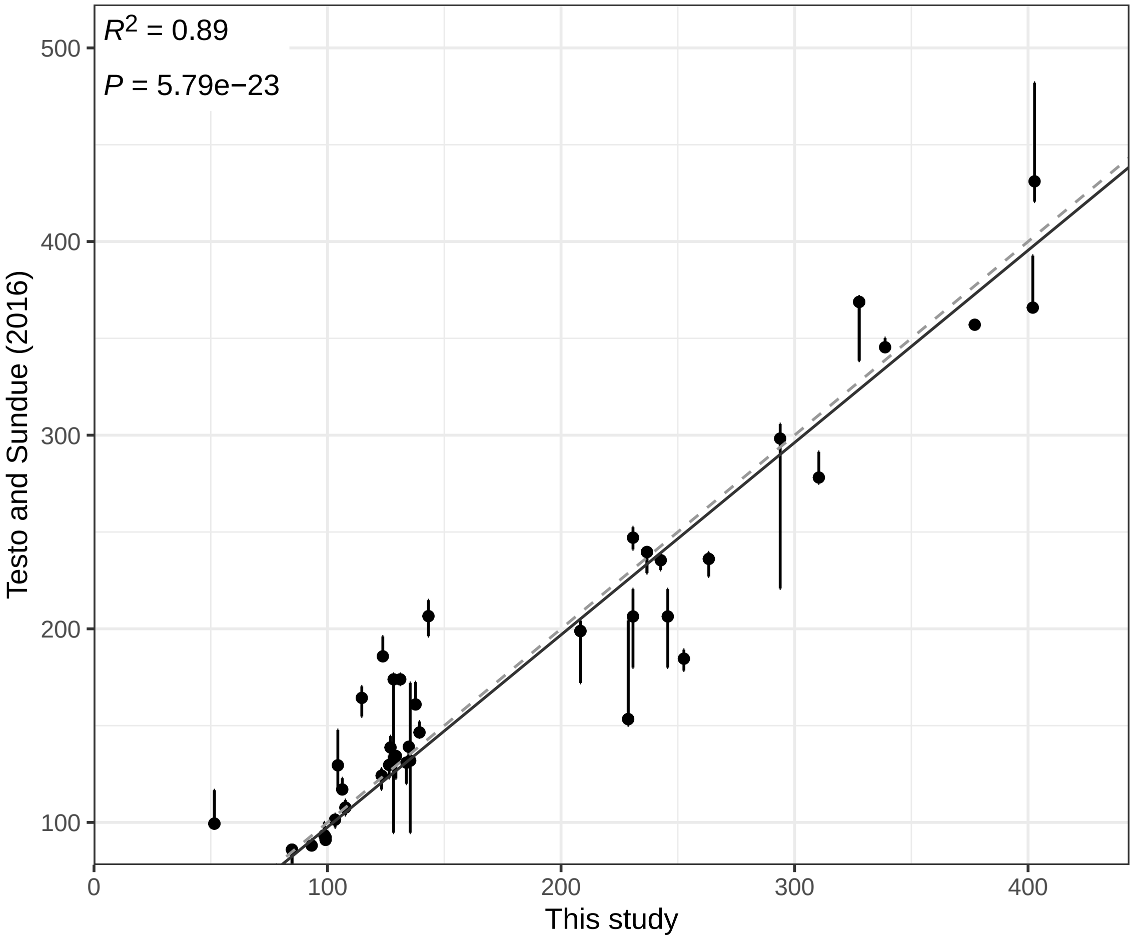


**Figure S10**. Comparison of stem age of fern families estimated by Testo and Sundue (2016) and those estimated using methodology (treePL) and phylogeny (FTOL) of the current study with the fossil constraints of Testo and Sundue (2016). All units in millions of years (Ma). For Testo and Sundue (2016), error bars indicate lower and upper 95% highest posterior density levels and point indicates median. For this study, point indicates best (most likely) estimate. Dashed indicates 1:1 relationship. Solid line indicates linear model fit.

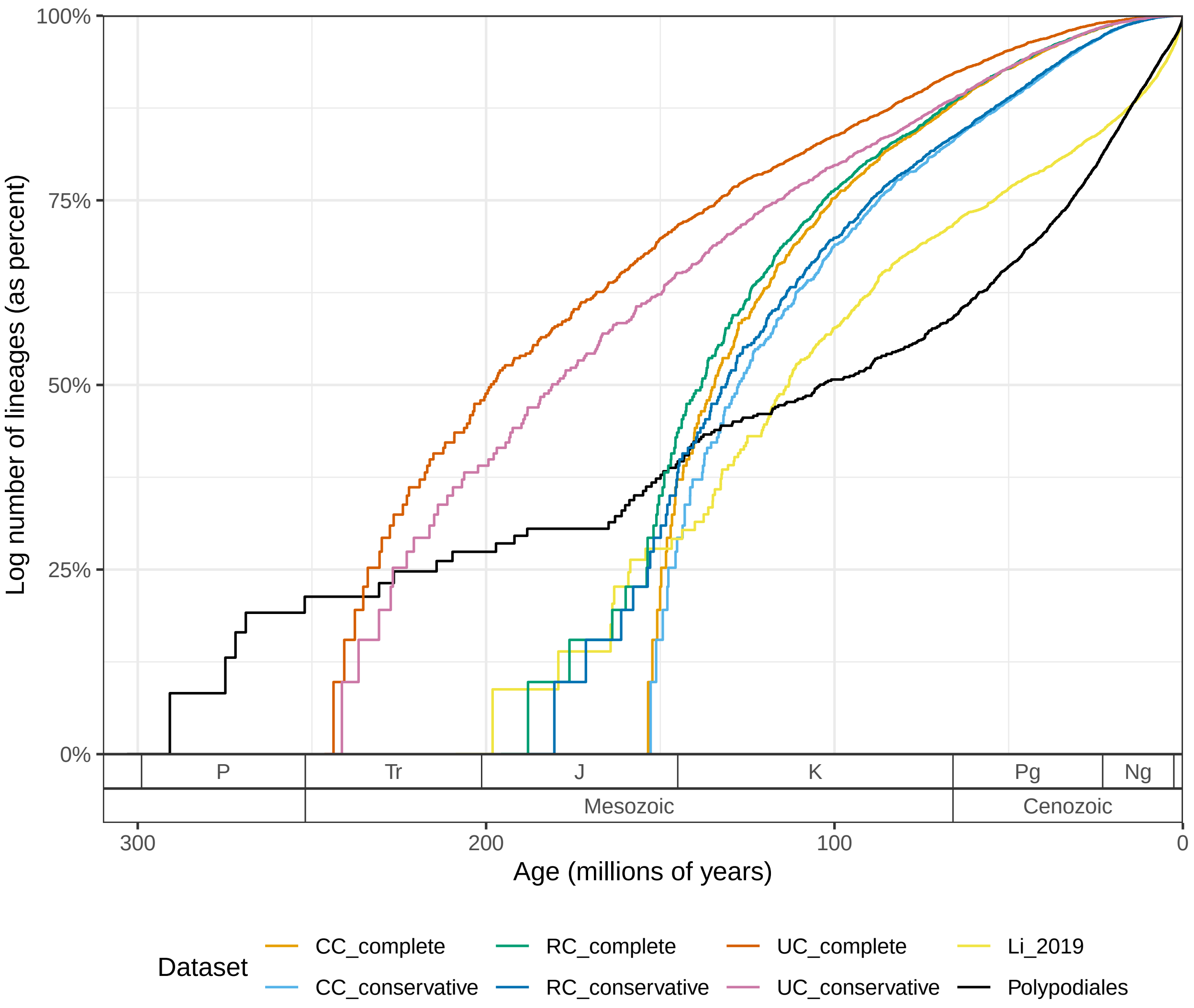


**Figure S11**. Lineage through time (LTT) plots for the fern order Polypodiales (black; this study) and angiosperms (colored lines; Li et al., 2019; Ramírez-Barahona et al., 2020). Angiosperm datasets labeled with "CC", "RC", or "UC" correspond to the six analysis schemes of Ramírez-Barahona et al. (2020); for details, see that paper. Log number of lineages shown as percent of total to enable comparison across trees.
