## Appendix A2 for "An Open and Continuously Updated Fern Tree of Life"

### Fossil Calibration List

These data are exported from the FernCal v1.0.1 database (https://github.com/fernphy/ferncal), which includes records of fossils to use as calibration points for molecular dating of ferns.

“NFos” is the unique ID of the record in FernCal.

Taxonomy of extant taxa follows pteridocat v1.0.0 database (https://github.com/fernphy/pteridocat).

For a summary of these data in CSV format, see Table S2.

### *Acrostichum intertrappeum*†

#### Calibration

- NFos: 35
- Minimum age: 66
- Node calibrated: stem *Acrostichum+Ceratopteris*

#### Fossil Age

- Lower limit of oldest stratigraphic age: 72.1
- Upper limit of oldest stratigraphic age: 66
- Stratigraphic age: Late Cretaceous (Maastrichtian)
- Reference time scale: ICS (v2017/02)

#### Fossil Identity

- Author: Bonde et Kumaran
- Full taxon name: *Acrostichum intertrappeum* Bonde et Kumaran
- Affinities (group): stem
- Affinities: *Acrostichum+Ceratopteris*
- Order (of clade calibrated): Polypodiales
- Family (of clade calibrated): Pteridaceae
- Notes on affinities: Characters of the fossil suggest its undoubted affinity with *Acrostichum*.
- Reference: Bonde and Kumaran (2002)

#### Fossil Locality

- Country: India
- Type locality: Nawargaon, District Wardha, Ma- harashtra, India, 21‡01PN, 78‡35PE
- Formation: Deccan Intertrappeans of India

#### Fossil Specimen

- Organs: Stem, petioles, roots
- Specimen (Holotype): N 201/98
- Collection: Department of Botany, Agharkar Research Institute, Pune, India

---

### *Acrostichum palaeoaureum*†

#### Calibration

- NFos: 36
- Minimum age: 23.03
- Node calibrated: stem *Acrostichum*

#### Fossil Age

- Lower limit of oldest stratigraphic age: 27.82
- Upper limit of oldest stratigraphic age: 23.03
- Stratigraphic age: Late Oligocene (Chattian)
- Reference time scale: ICS (v2017/02)
- Notes on age: Ash-IV is a 3–7-m-thick, basin-wide unit that occurs stratigraphically above the basalt and is dated by 40Ar/39Ar at 27.36 ± 0.11 Ma.

#### Fossil Identity

- Author: Beauchamp, Lemoigne et Petrescu
- Full taxon name: *Acrostichum palaeoaureum* Beauchamp, Lemoigne et Petrescu
- Affinities (group): stem
- Affinities: *Acrostichum*
- Order (of clade calibrated): Polypodiales
- Family (of clade calibrated): Pteridaceae
- Reference: García-Massinni et al. (2006)

#### Fossil Locality

- Country: Etiopia
- Type locality: Chilga, ca. 60 km west-southwest of Gondar on the northwestern Ethiopian Plateau, along the slopes of the Magargaria River (12°30931.30N, 37°6957.30E) and along the Guang River (12°30943.440N, 37°7924.180E)
- Formation: Volcanic ash deposits within Ash-IV in the Guang River section and lignitic deposits within Ash- IV in the Magargaria River

#### Fossil Specimen

- Organs: Sterile foliage, stems, spores
- Specimen (Holotype): CH-55 6 (fig. 2A), CH-55 4 (fig. 2B), CH-55 2 (fig. 2C), CH-79 21 A (fig. 2D), CH-79 21 B (fig. 2E), and CH-79 21 A-B #1 (fig. 2F–2H)
- Collection: National Museum of Ethiopia, Addis Abada

---

### *Aglaomorpha heraclea*

#### Calibration

- NFos: 4
- Minimum age: 7.246
- Node calibrated: crown *Aglaomorpha*

#### Fossil Age

- Lower limit of oldest stratigraphic age: 11.63

- Upper limit of oldest stratigraphic age: 7.246
- Stratigraphic age: Upper Miocene (Messinian-Tortonian)
- Reference time scale: ICS (v2017/02)

#### Fossil Identity

- Author: (Kunze) Copel
- Full taxon name: *Aglaomorpha heraclea* (Kunze) Copel
- Affinities (group): crown
- Affinities: *Aglaomorpha*
- Order (of clade calibrated): Polypodiales
- Family (of clade calibrated): Polypodiaceae
- Notes on affinities: Uffelen (1991) mentions an assignment to *Drynaria* and to *Aglaomorpha heraclea*; conservative assignment to crown *Aglaomorpha*.
- Reference: Uffelen (1991)

#### Fossil Locality

- Country: Sumatra
- Type locality: Palembang Province

#### Fossil Specimen

- Organs: Fertile foliage
- Collection: Natural History Museum

---

### *Asplenium sanshuiense*†

#### Calibration

- NFos: 37
- Minimum age: 47.8
- Node calibrated: stem *Asplenium*

#### Fossil Age

- Lower limit of oldest stratigraphic age: 56
- Upper limit of oldest stratigraphic age: 47.8
- Stratigraphic age: early Eocene (Ypresian)
- Reference time scale: ICS (v2017/02)

#### Fossil Identity

- Author: Xu et Jin
- Full taxon name: *Asplenium sanshuiense* Xu et Jin
- Affinities (group): stem

- Affinities: *Asplenium*
- Order (of clade calibrated): Polypodiales
- Family (of clade calibrated): Aspleniaceae
- Notes on affinities: The frond shape, the rachis structure and the venation characteristic of ultimate segment in fossil specimens mostly resemble the extant species of *Asplenium*.
- Reference: Xu et al. (2017b)

#### Fossil Locality

- Country: China
- Type locality: Sanshui Basin, Guangdong Province
- Formation: Huachong Fm

#### Fossil Specimen

- Organs: Sterile fronds
- Specimen (Holotype): SSHC-011 (holotype)
- Collection: Museum of Biology, Sun Yet-sen University, Guangzhou, China

---

### *Athyrium cretaceum*†

#### Calibration

- NFos: 119
- Minimum age: 129.4
- Node calibrated: stem *Athyrium* s.s.

#### Fossil Age

- Lower limit of oldest stratigraphic age: 145
- Upper limit of oldest stratigraphic age: 129.4
- Stratigraphic age: Neocomian (Barresian to Hauterivian)
- Reference time scale: ICS (v2017/02)
- Notes on age: The age is Aptian-Neocomian based on stratigraphic correlations.

#### Fossil Identity

- Author: Chen et Meng
- Full taxon name: *Athyrium cretaceum* Chen et Meng
- Affinities (group): stem
- Affinities: *Athyrium* s.s.
- Order (of clade calibrated): Polypodiales
- Family (of clade calibrated): Athyriaceae
- Notes on affinities: The fossil is compared to species of *Athyrium* s.s. (not sensu Pteridophyte Phylogeny Group I, 2016). Taken as-is to calibrate *Athyrium* s.s.
- Reference: Chen et al. (1997)

**Fossil Locality**

- Country: China
- Type locality: Fuxin Basin and Tiefa Basin, Liaoning province; Huolinnhe Basin, Inner Mongolia
- Formation: Fuxin and Xiaominganbei Formations; Huolinhe Formation

**Fossil Specimen**

- Organs: Fertile fronds, sporangia, spores
- Specimen (Holotype): Plates 1 and 2
- Collection: China University of Geosciences

---

### *Cibotiocaulis tateiwai*†

**Calibration**

- NFos: 46
- Minimum age: 129.4
- Node calibrated: stem Cyatheaceae

**Fossil Age**

- Lower limit of oldest stratigraphic age: 133.9
- Upper limit of oldest stratigraphic age: 129.4
- Stratigraphic age: Hauterivian
- Reference time scale: ICS (v2017/02)

**Fossil Identity**

- Author: (Ogura) Ogura
- Full taxon name: *Cibotiocaulis tateiwai* (Ogura) Ogura
- Affinities (group): stem
- Affinities: Cyatheaceae
- Order (of clade calibrated): Cyatheales
- Family (of clade calibrated): Cyatheaceae
- Notes on affinities: Affinities within the Cyatheaceae. According to Stockey and Rothwell (2004) and Lantz et al. (1999), the fossil can be attributed to stem Cyatheaceae
- Reference: Nishida (1989)

**Fossil Locality**

- Country: Korea
- Type locality: North Kyong Sang Province
- Formation: Not provided

#### Fossil Specimen

- Organs: Stem
- Specimen (Holotype): Specimen 5 (holotype)
- Collection: University Museum of the University of Tokyo

---

### *Cibotium oregonense*†

#### Calibration

- NFos: 9
- Minimum age: 33.9
- Node calibrated: crown Cibotiaceae

#### Fossil Age

- Lower limit of oldest stratigraphic age: 37.8
- Upper limit of oldest stratigraphic age: 33.9
- Stratigraphic age: Upper Eocene (?)
- Reference time scale: ICS (v2017/02)
- Notes on age: Terrestrial sedimentary rocks in the region of Medford are all Upper Eocene.

#### Fossil Identity

- Author: Barrington
- Full taxon name: *Cibotium oregonense* Barrington
- Affinities (group): crown
- Affinities: Cibotiaceae
- Order (of clade calibrated): Cyatheales
- Family (of clade calibrated): Cibotiaceae
- Notes on affinities: A placement within the Cibotiaceae is suggested. According to Stockey and Rothwell (2004), the fossil nests within extant Cibotiaceae and Metaxyaceae. Lantz et al. (1999) place it separate from extant *Cibotium barometz*, in a clade including *Metaxya rostrata*.
- Reference: Barrington (1983)

#### Fossil Locality

- Country: North America
- Type locality: Unknown, Medford, Oregon
- Formation: Uncertain

#### Fossil Specimen

- Organs: Stem and petioles
- Specimen (Holotype): 60674

- Collection: Paleobotanical Collections of the Botanical Museum, Harvard University

---

### *Coniopteris lobata*†

#### Calibration

- NFos: 56
- Minimum age: 174.1
- Node calibrated: stem Cyatheales

#### Fossil Age

- Lower limit of oldest stratigraphic age: 201.3
- Upper limit of oldest stratigraphic age: 174.1
- Stratigraphic age: Lower Jurassic
- Reference time scale: ICS (v2017/02)

#### Fossil Identity

- Author: (Oldham and Morris) Halle
- Full taxon name: *Coniopteris lobata* (Oldham and Morris) Halle
- Affinities (group): stem
- Affinities: Cyatheales
- Order (of clade calibrated): Cyatheales
- Reference: Rees and Cleal (2004)

#### Fossil Locality

- Country: Antartica
- Type locality: Hope Bay and Botany Bay
- Formation: Not provided

#### Fossil Specimen

- Organs: Sterile and fertile foliage
- Specimen (Holotype): D.8911.1B, D.8916B, D.8047.4A, D.8947.4B, D.8947.11A, D.8947.17B, D.8958.10A; 8003
- Collection: The Natural History Museum, London

---

### *Culcita emberi*†

#### Calibration

- NFos: 10

- Minimum age: 13.82
- Node calibrated: crown Culcitaceae

**Fossil Age**

- Lower limit of oldest stratigraphic age: 15.97
- Upper limit of oldest stratigraphic age: 13.82
- Stratigraphic age: Middle Miocene
- Reference time scale: ICS (v2017/02)

**Fossil Identity**

- Author: Pinson, Manchester et Sessa
- Full taxon name: *Culcita emberi* Pinson, Manchester et Sessa
- Affinities (group): crown
- Affinities: Culcitaceae
- Order (of clade calibrated): Cyatheales
- Family (of clade calibrated): Culcitaceae
- Notes on affinities: The oblique, complete annulus surely puts it within the Cyatheales.
- Reference: Pinson et al. (2018)

**Fossil Locality**

- Country: North America
- Type locality: Clarkia Beds
- Formation: Emerald Creek locality (Univerdity of Idaho P-37)

**Fossil Specimen**

- Organs: Fertile foliage, sporangia, spores
- Specimen (Holotype): UF 18630-61360 (holotype)
- Collection: Paleobotanical Collection of the Florida Museum of Natural History, University of Florida

---

### *Cyatheaceae sp. indet.*†

**Calibration**

- NFos: 20
- Minimum age: 7.246
- Node calibrated: crown Cyathea

**Fossil Age**

- Lower limit of oldest stratigraphic age: 11.63
- Upper limit of oldest stratigraphic age: 7.246

- Stratigraphic age: Upper Miocene (Messinian-Tortonian)
- Reference time scale: ICS (v2017/02)

#### Fossil Identity

- Author: Graham
- Full taxon name: Cyatheaceae *sp. indet.* Graham
- Affinities (group): crown
- Affinities: *Cyathea*
- Order (of clade calibrated): Cyatheales
- Family (of clade calibrated): Cyatheaceae
- Notes on affinities: Most similarity with *Sphaeropteris myosuroides* and *Trichipteris mexicana*. The two are synonyms of New World *Cyathea*.
- Reference: Graham (1979)

#### Fossil Locality

- Country: Mexico
- Type locality: Coatzacoalcos
- Formation: Paraje Solo Fm.

#### Fossil Specimen

- Organs: Spores
- Specimen (Holotype): Slide 10-1, England Slide finder coordinates C-45

---

### *Cyclosorus scutum*†

#### Calibration

- NFos: 59
- Minimum age: 33.9
- Node calibrated: stem *Cyclosorus*

#### Fossil Age

- Lower limit of oldest stratigraphic age: 56
- Upper limit of oldest stratigraphic age: 33.9
- Stratigraphic age: Eocene
- Reference time scale: ICS (v2017/02)
- Notes on age: The fossils of *Cyclosorus*, described here, were collected from the coal-bearing deposits of the Changchang Formation assigned to an Eocene age according to the published paleobotanical (megafossil) and palynological data.

#### Fossil Identity

- Author: Naugolnykh, Wang, Han et Jin
- Full taxon name: *Cyclosorus scutum* Naugolnykh, Wang, Han et Jin
- Affinities (group): stem
- Affinities: Cyclosorus
- Order (of clade calibrated): Polypodiales
- Family (of clade calibrated): Thelypteridaceae
- Notes on affinities: The material studied can be classified into *Cyclosorus* (Thelypteridaceae) with certainty, despite the exact attribution of *C. scutum* at the subgeneric level remaining unclear.
- Reference: Naugolnykh et al. (2016)

#### Fossil Locality

- Country: China
- Type locality: Jiazi Town, the City of Qiongshan, northeastern part of the Hainan Island, China
- Formation: Changchang Formation

#### Fossil Specimen

- Organs: Sterile and fertile fronds, sporangia, spores
- Specimen (Holotype): Holotype: CCJ 266A-f-1
- Collection: Museum of Biology, Sun Yet-sen University, Guangzhou, China

---

### *Dicksonia mariopteris*†

#### Calibration

- NFos: 21
- Minimum age: 168.3
- Node calibrated: crown Cyatheaceae+Dicksoniaceae+Cibotiaceae+Metaxyaceae

#### Fossil Age

- Lower limit of oldest stratigraphic age: 170.3
- Upper limit of oldest stratigraphic age: 168.3
- Stratigraphic age: Bajocian
- Reference time scale: ICS (v2017/02)

#### Fossil Identity

- Author: Wilson et Yates
- Full taxon name: *Dicksonia mariopteris* Wilson et Yates
- Affinities (group): crown
- Affinities: Cyatheaceae+Dicksoniaceae+Cibotiaceae+Metaxyaceae

- Order (of clade calibrated): Cyatheales
- Notes on affinities: The author assigned these fossils to the Dicksoniae (incl. *Cibotium*, *Culcita* subgenus *Calochlaena*, *Cystodium*, *Dicksonia*). Thus the assignment to the Dicksoniaceae is not justified. Hence, we conservatively treat the fossil as member of the core tree ferns.
- Reference: van Konijnenburg-van Cittert (1989)

#### Fossil Locality

- Country: UK
- Type locality: Roseberry Topping, Hasty Bank
- Formation: Saltwick Fm.

#### Fossil Specimen

- Organs: Spores
- Specimen (Holotype): v.32252 $1 (type specimen), v.32252, v.56792, v.60980A
- Collection: British Museum (Natural History)

---

### *Dryopteris guyottii*†

#### Calibration

- NFos: 11
- Minimum age: 33.9
- Node calibrated: stem *Dryopteris*

#### Fossil Age

- Lower limit of oldest stratigraphic age: 37.8
- Upper limit of oldest stratigraphic age: 33.9
- Stratigraphic age: Priabonian
- Reference time scale: ICS (v2017/02)
- Notes on age: Radioisotopic age of 34.07–36.7 M (40Ar/39Ar radiometry).

#### Fossil Identity

- Author: (Lesquereux) MacGinitie
- Full taxon name: *Dryopteris guyottii* (Lesquereux) MacGinitie
- Affinities (group): stem
- Affinities: *Dryopteris*
- Order (of clade calibrated): Polypodiales
- Family (of clade calibrated): Dryopteridaceae
- Notes on affinities: Affinities with *Dryopteris cristata* and *D. floridana*.
- Reference: MacGinitie (1953)

**Fossil Locality**

- Country: United States
- Type locality: Florissant and Florissant Fossil Beds National Monument, Colorado
- Formation: Florissant Fm.

**Fossil Specimen**

- Organs: Fertile foliage
- Specimen (Holotype): Syntypes: 1613, 1614
- Collection: US National Museum

---

### *Dryopterites beishanensis*†

**Calibration**

- NFos: 140
- Minimum age: 129.4
- Node calibrated: stem Dryopteridaceae

**Fossil Age**

- Lower limit of oldest stratigraphic age: 132.9
- Upper limit of oldest stratigraphic age: 129.4
- Stratigraphic age: Early Cretaceous (Hauterivian-Barremian)
- Reference time scale: ICS (v2017/02)

**Fossil Identity**

- Author: Ren et Sun
- Full taxon name: *Dryopterites beishanensis* Ren et Sun
- Affinities (group): stem
- Affinities: Dryopteridaceae
- Order (of clade calibrated): Polypodiales
- Family (of clade calibrated): Dryopteridaceae
- Reference: Ren et al. (2022)

**Fossil Locality**

- Country: China
- Type locality: Zhongkouzi Basin, Beishan area, Northwest China
- Formation: Chijinbao Fm.

**Fossil Specimen**

- Organs: Fertile foliage, sporangia, spores

- Specimen (Holotype): 19-k-2-337
- Collection: Palaeontology Laboratory of the School of Earth Sciences, Lanzhou University, Lanzhou, China

---

### *Elaphoglossum miocenicum*†

#### Calibration

- NFos: 63
- Minimum age: 20.44
- Node calibrated: stem *Elaphoglossum*

#### Fossil Age

- Lower limit of oldest stratigraphic age: 23.03
- Upper limit of oldest stratigraphic age: 20.44
- Stratigraphic age: Early Miocene (Burdigalian-Aquitanian)
- Reference time scale: ICS (v2017/02)
- Notes on age: The amber has been dated as early Miocene, 20 to 15 Myr old, and was exuded by resin-bearing species of Hymenaea in the Fabaceae.

#### Fossil Identity

- Author: Lóriga, A.R.Schmidt, R.C.Moran, K.Feldberg, H.Schneid. et Heinrichs
- Full taxon name: *Elaphoglossum miocenicum* Lóriga, A.R.Schmidt, R.C.Moran, K.Feldberg, H.Schneid. et Heinrichs
- Affinities (group): stem
- Affinities: *Elaphoglossum*
- Order (of clade calibrated): Polypodiales
- Family (of clade calibrated): Dryopteridaceae
- Reference: Lóriga et al. (2014)

#### Fossil Locality

- Country: Dominincan Republic
- Type locality: Dominican Republic, Santiago area
- Formation: Not provided

#### Fossil Specimen

- Organs: Fertile foliage, petiole, sporangia, spores
- Specimen (Holotype): USNM 414283
- Collection: amber collection of the U. S. National Museum of Natural History at the Smithsonian Institution

---

### *Eophyllophyton bellum*†

#### Calibration

- NFos: 131
- Minimum age: 407.6
- Node calibrated: stem Euphyllophytes

#### Fossil Age

- Lower limit of oldest stratigraphic age: 410.8
- Upper limit of oldest stratigraphic age: 407.6
- Stratigraphic age: Early Devonian (Pragian)
- Reference time scale: ICS (v2017/02)

#### Fossil Identity

- Author: Hao
- Full taxon name: *Eophyllophyton bellum* Hao
- Affinities (group): stem
- Affinities: Euphyllophytes
- Reference: Hao and Xue (2013)

#### Fossil Locality

- Country: China
- Type locality: Zhichang Village
- Formation: Posongchong Fm.

#### Fossil Specimen

- Organs: Rhizomes, stems, leaves, sporangia
- Specimen (Holotype): Plate 4, fig 9 and Plate 6, fig 8 (from Hao & Beck 1993); Plate III, fig 4 and Plate II fig 11 (from HAo 1988)
- Collection: Not provided

---

### *Equisetum cf. oppositum*†

#### Calibration

- NFos: 27
- Minimum age: 56
- Node calibrated: crown *Equisetum* subgen. *Equisetum*

#### Fossil Age

- Lower limit of oldest stratigraphic age: 66
- Upper limit of oldest stratigraphic age: 56
- Stratigraphic age: Paleocene-Eocene
- Reference time scale: ICS (v2017/02)
- Notes on age: Combined with plant compressions, charophytes, ostracodes, and palynological fossils from the Niubao Formation and overlying Dingqinghu Formation indicate that the age of measured section is the Paleocene-Eocene.

#### Fossil Identity

- Author: Ma, Su et Zhang
- Full taxon name: *Equisetum* cf. *oppositum* Ma, Su et Zhang
- Affinities (group): crown
- Affinities: *Equisetum* subgen. *Equisetum*
- Order (of clade calibrated): Equisetales
- Family (of clade calibrated): Equisetaceae
- Notes on affinities: Assigned to subgenus *Equisetum* based on morphological similarity, and absence of tubers in subgenus *Hippochaete*.
- Reference: Yu et al. (2020)

#### Fossil Locality

- Country: China
- Type locality: Nima County, the Tibet Autonomous Region, southwestern China
- Formation: Niubao Fm

#### Fossil Specimen

- Organs: Rhizomes, tubers
- Specimen (Holotype): XZNM-2013LDGSW-EN-01; XZNM-2013DGSW-EN-02
- Collection: School of Earth Science, Lanzhou University

---

### *Equisetum clarnoi*†

#### Calibration

- NFos: 28
- Minimum age: 41.2
- Node calibrated: crown *Equisetum* subgen. *Hippochaete*

#### Fossil Age

- Lower limit of oldest stratigraphic age: 47.8
- Upper limit of oldest stratigraphic age: 41.2

- Stratigraphic age: Middle Eocene (Lutetian)
- Reference time scale: ICS (v2017/02)

**Fossil Identity**

- Author: Brown
- Full taxon name: *Equisetum clarnoi* Brown
- Affinities (group): crown
- Affinities: *Equisetum* subgen. *Hippochaete*
- Order (of clade calibrated): Equisetales
- Family (of clade calibrated): Equisetaceae
- Notes on affinities: Christenhusz et al. (2021) assigns this fossil to subgenus *Hippochaete*; agreement with Elgorriaga et al. (2018).
- Reference: Brown (1975)

**Fossil Locality**

- Country: North America
- Type locality: NW'/4, Sec. 27, T9S, R18E; 91'/ miles E, 1 mile N of Ashwood, Jefferson Co., Oregon
- Formation: Clarno Chert

**Fossil Specimen**

- Organs: Stem and roots
- Specimen (Holotype): PB #292
- Collection: University of Montana Paleontological Collectio

---

### *Equisetum laterale*†

**Calibration**

- NFos: 136
- Minimum age: 174.1
- Node calibrated: stem *Equisetum* subgen. *Paramochaete*

**Fossil Age**

- Lower limit of oldest stratigraphic age: 201.3
- Upper limit of oldest stratigraphic age: 174.1
- Stratigraphic age: Lower Jurassic
- Reference time scale: ICS (v2017/02)

**Fossil Identity**

- Author: Phillips emend. Gould

- Full taxon name: *Equisetum laterale* Phillips emend. Gould
- Affinities (group): stem
- Affinities: *Equisetum* subgen. *Paramochaete*
- Order (of clade calibrated): Equisetales
- Family (of clade calibrated): Equisetaceae
- Notes on affinities: Christenhusz et al. (2021) assign this fossil to subgen. *Hippochaete* in agreement with Elgorriaga et al. (2018).
- Reference: Rees and Cleal (2004)

**Fossil Locality**

- Country: Antartica
- Type locality: Hope Bay and Botany Bay
- Formation: Not provided

**Fossil Specimen**

- Organs: Stems, leaf sheaths
- Specimen (Holotype): D.34.7, D.37.1a, D.37.1c, D.37.2, D.37.7, D.37.9, D.39.6; D.8878, D.8932.3A, D.8935.2, D.8935.5, D.8935.8B, D.8936.3, D.8955.1, D.8965.14A, D.8965.25A, D.8965.32B, D.8974.2, D.8975.1c, D.8975.4B
- Collection: The Natural History Museum, London

---

### *Gleichenia chaloneri*†

**Calibration**

- NFos: 67
- Minimum age: 100.5
- Node calibrated: stem *Gleichenia*

**Fossil Age**

- Lower limit of oldest stratigraphic age: 113
- Upper limit of oldest stratigraphic age: 100.5
- Stratigraphic age: lower Albian
- Reference time scale: ICS (v2017/02)

**Fossil Identity**

- Author: Herendeen et J.Skog
- Full taxon name: *Gleichenia chaloneri* Herendeen et J.Skog
- Affinities (group): stem
- Affinities: *Gleichenia*
- Order (of clade calibrated): Gleicheniales
- Family (of clade calibrated): Gleicheniaceae

- Notes on affinities: Cladistic analysis places the fossil sister to *Gleichenia glauca*, clearly segregated from other genera; however, a single extant species of Gleichenia does not allow to ascertain whether the fossil lays within the crown group so we conservatively place the fossil as stem.
- Reference: Herendeen and Skog (1998)

**Fossil Locality**

- Country: UK
- Type locality: Munday'sHill Quarry,LeightonBuzzard,Bedfordshire, England

**Fossil Specimen**

- Organs: Fertile foliage, sporagia
- Specimen (Holotype): Specimen PP45078
- Collection: paleobotanical collections of the Department of Geology, The Field Museum (PP)

---

### *Glomerisporites pupus*†

**Calibration**

- NFos: 38
- Minimum age: 83.6
- Node calibrated: stem *Azolla*

**Fossil Age**

- Lower limit of oldest stratigraphic age: 86.3
- Upper limit of oldest stratigraphic age: 83.6
- Stratigraphic age: Santonian to (?) early Campanian
- Reference time scale: ICS (v2017/02)
- Notes on age: Originally dated as (middle) Senonian, the more precise determination of Santonian to possibly early Campanian is now generally accepted.

**Fossil Identity**

- Author: (Dijkstra) Potonié
- Full taxon name: *Glomerisporites pupus* (Dijkstra) Potonié
- Affinities (group): stem
- Affinities: *Azolla*
- Order (of clade calibrated): Salviniales
- Family (of clade calibrated): Salviniaceae
- Notes on affinities: Re-analysis of Dijkstra (1949) collection and spore-wall morphology confirm evolutionary links to several taxa within the Salviniales and other fossils possibly related to the group.
- Reference: Batten et al. (1998)

**Fossil Locality**

- Country: Europe
- Type locality: Epen, Vaals, and near Simpelveld
- Formation: Aachen Fm.

**Fossil Specimen**

- Organs: Megaspore
- Specimen (Holotype): SEM micrographs depicted in figure (all from Dijkstra's slide collection)
- Collection: Not provided

---

### *Goniophlebium macrosorum*†

**Calibration**

- NFos: 70
- Minimum age: 15.97
- Node calibrated: stem *Goniophlebium*

**Fossil Age**

- Lower limit of oldest stratigraphic age: 20.44
- Upper limit of oldest stratigraphic age: 15.97
- Stratigraphic age: Miocene (Langhian-Burdigalian)
- Absolute age: 15.2–16.5 Ma
- Reference time scale: ICS (v2017/02)
- Notes on age: According to a recent magnetostratigraphic study, the sedimentary succession was assigned to Middle Miocene (15.2–16.5 Ma).

**Fossil Identity**

- Author: Xu et Zhou
- Full taxon name: *Goniophlebium macrosorum* Xu et Zhou
- Affinities (group): stem
- Affinities: *Goniophlebium*
- Order (of clade calibrated): Polypodiales
- Family (of clade calibrated): Polypodiaceae
- Reference: Xu et al. (2017a)

**Fossil Locality**

- Country: China
- Type locality: Dashidong village, the Wenshan Basin, southeastern Yunnan, China
- Formation: Xiaolongtan Fm

#### Fossil Specimen

- Organs: Sterile and fertile foliage
- Specimen (Holotype): KUN PC2016DMS0368
- Collection: Herbarium of Kunming Institute of Botany, Chinese Academy of Sciences (KUN)

---

### *Heinrichsia cheilanthoides*†

#### Calibration

- NFos: 117
- Minimum age: 100.5
- Node calibrated: crown Pteridaceae

#### Fossil Age

- Lower limit of oldest stratigraphic age: 113
- Upper limit of oldest stratigraphic age: 100.5
- Stratigraphic age: Late Albian-early Cenomanian
- Reference time scale: ICS (v2017/02)
- Notes on age: Biostratigraphic studies (Cruickshank & Ko, 2003) and U-Pb dating of zircons indicate a late Albian to earliest Cenomanian age of the Burmese amber, with a minimum age of 98 Ma.

#### Fossil Identity

- Author: Regalado, A.R.Schmidt, M.Krings et H.Schneid.
- Full taxon name: *Heinrichsia cheilanthoides* Regalado, A.R.Schmidt, M.Krings et H.Schneid.
- Affinities (group): crown
- Affinities: Pteridaceae
- Order (of clade calibrated): Polypodiales
- Family (of clade calibrated): Pteridaceae
- Notes on affinities: The fossil was assigned to the order Polypodiales based on the presence of a vertical annulus interrupted by the sporangium stalk. The presence of marginal sori protected by pseudoindusia therefore is strong support of the affinities of the fossil to the Pteridaceae.
- Reference: Regalado et al. (2019)

#### Fossil Locality

- Country: Myanmar
- Type locality: Amber mines near Tanai, Ledo Road, 105 km northwest of Myitkyina, Kachin State
- Formation: Burmese amber (Burmite)

#### Fossil Specimen

- Organs: Fertile foliage, sporangia, spores
- Specimen (Holotype): GZG.BST.21977
- Collection: Collections of the Geoscience Centre (GZG) at the University of Göttingen

---

### *Hymenophyllum axsmithii*†

#### Calibration

- NFos: 148
- Minimum age: 47.8
- Node calibrated: crown *Hymenophyllum*

#### Fossil Age

- Lower limit of oldest stratigraphic age: 56
- Upper limit of oldest stratigraphic age: 47.8
- Stratigraphic age: Early Eocene (Ypresian)
- Absolute age: 49.4 + 0.5 Ma
- Reference time scale: ICS (v2017/02)
- Notes on age: An unpublished 40Ar-39Ar date of 49.4+0.5 Ma also places the Republic flora as latest early Eocene

#### Fossil Identity

- Author: Pigg, Greenwood, Sundue et DeVore
- Full taxon name: *Hymenophyllum axsmithii* Pigg, Greenwood, Sundue et DeVore
- Affinities (group): crown
- Affinities: *Hymenophyllum*
- Order (of clade calibrated): Hymenophyllales
- Family (of clade calibrated): Hymenophyllaceae
- Notes on affinities: Affinities with *Hymenophyllum* subgen. *Sphaerocionium*.
- Reference: Pigg et al. (2021)

#### Fossil Locality

- Country: United States
- Type locality: Boot Hill, locality B4131, Republic, Washington (Holotype)
- Formation: Klondike Mountain fm.

#### Fossil Specimen

- Organs: Fertile foliage, sporangia, spores (?)
- Specimen (Holotype): SR 05-15-13
- Collection: Burke Museum of Natural History and Culture

---

### *Hymenophyllum iwatsukii*†

#### Calibration

- NFos: 72
- Minimum age: 113
- Node calibrated: stem *Hymenophyllum*

#### Fossil Age

- Lower limit of oldest stratigraphic age: 125
- Upper limit of oldest stratigraphic age: 113
- Stratigraphic age: Aptian-Albian
- Reference time scale: ICS (v2017/02)
- Notes on age: Aptian-Albian age based on stratigraphic correlations and palynomorphs recovered from the plant localities.

#### Fossil Identity

- Author: Herrera, R.C.Moran, Shi, Ichinnorov, Takahashi, Crane et Herendeen
- Full taxon name: *Hymenophyllum iwatsukii* Herrera, R.C.Moran, Shi, Ichinnorov, Takahashi, Crane et Herendeen
- Affinities (group): stem
- Affinities: *Hymenophyllum*
- Order (of clade calibrated): Hymenophyllales
- Family (of clade calibrated): Hymenophyllaceae
- Notes on affinities: Assigned to the Hymnophyllaceae based on a combination of vegetative and reproductive characters.
- Reference: Herrera et al. (2017)

#### Fossil Locality

- Country: Mongolia
- Type locality: Tevshiin Govi (45°58'54" N, 106°07'12" E) and Tugrug (46°53'05" N, 108°03'38" E) coal mines, central Mongolia
- Formation: Tevshiin Govi and Khukhteeg Fm

#### Fossil Specimen

- Organs: Fertile fronds, sporangia, spores
- Specimen (Holotype): PP56496
- Collection: Paleobotanical Collections, Department of Geology, The Field Museum, Chicago, Illinois, USA

---

### *Klukia exilis*†

#### Calibration

- NFos: 100
- Minimum age: 163.5
- Node calibrated: stem Schizaeaceae

#### Fossil Age

- Lower limit of oldest stratigraphic age: 174.1
- Upper limit of oldest stratigraphic age: 163.5
- Stratigraphic age: Middle Jurassic
- Reference time scale: ICS (v2017/02)

#### Fossil Identity

- Author: (Phillips) Raciborski
- Full taxon name: *Klukia exilis* (Phillips) Raciborski
- Affinities (group): stem
- Affinities: Schizaeaceae
- Order (of clade calibrated): Schizaeales
- Family (of clade calibrated): Schizaeaceae
- Notes on affinities: Comparing the Jurassic *Klukia exilis* and *Stachypteris spicans* spores with Recent spores, one may say that they are similar but not strictly comparable, especially their greatly reduced proximal sculpture that has not been seen in the Recent spores. Within the Schizaeaceae, however, they certainly resemble most closely reticulate *Lygodium* spores (van Konijnenburg-van Cittert, 1989); therefore these may mark the split between *Lygodium* and the *Anemia*/*Schizaea* clade.
- Reference: Harris (1961)

#### Fossil Locality

- Country: UK
- Type locality: Cloughton Wyke and CaytonBay
- Formation: Middle Deltaic Gristhorpe Bed

#### Fossil Specimen

- Organs: Sterile and fertile fronds, sporangia, spores
- Specimen (Holotype): V.26903, V.25840, V.31973, V.31973a; Leckenby Coll., 91
- Collection: British Museum (Natural History) and Sedgwick Museum, Cambridge

---

### *Krameropteris resinatus*†

#### Calibration

- NFos: 92
- Minimum age: 100.5
- Node calibrated: stem *Monachosorum*

#### Fossil Age

- Lower limit of oldest stratigraphic age: 113
- Upper limit of oldest stratigraphic age: 100.5
- Stratigraphic age: late Albian
- Reference time scale: ICS (v2017/02)
- Notes on age: Biostratigraphic studies suggested a late Albian age of the amber-bearing sediment, hence the inclusions have a late Early Cretaceous age, with a minimum age of 98 million years (earliest Cenomanian, early Late Cretaceous) that is based on recent U-Pb dating of zircons.

#### Fossil Identity

- Author: H.Schneid., A.R.Schmidt et Heinrichs
- Full taxon name: *Krameropteris resinatus* H.Schneid., A.R.Schmidt et Heinrichs
- Affinities (group): stem
- Affinities: *Monachosorum*
- Order (of clade calibrated): Polypodiales
- Family (of clade calibrated): Dennstaedtiaceae
- Notes on affinities: *Monachosorum* species resemble the fossil in the formation of trilete spores, presence of exindusiate sori and a simultaneous maturity of sporangia. Given the uncertainty of character state evolution, the fossil may not be a member of the stem of *Monachosorum* as considered here but actually better placed at the branch leading to the split between *Monachosorum* and the HYP-clade. However, this appears to be less likely given the character state reconstructions obtained.
- Reference: Schneider et al. (2016)

#### Fossil Locality

- Country: Myanmar
- Type locality: Amber mines near Tanai, about 105 km north of Myitkyina in Kachin State
- Formation: Burmese amber (Burmite)

#### Fossil Specimen

- Organs: Fertile foliage, sporangia, spores
- Specimen (Holotype): Holotype: AMNH Bu-ASJH-3.
- Collection: American Museum of Natural History, New York

### *Kuylisporites mirabilis*†

#### Calibration

- NFos: 42
- Minimum age: 93.9
- Node calibrated: stem *Cyathea*

#### Fossil Age

- Lower limit of oldest stratigraphic age: 100.5
- Upper limit of oldest stratigraphic age: 93.9
- Stratigraphic age: Cenomanian
- Reference time scale: ICS (v2017/02)

#### Fossil Identity

- Full taxon name: *Kuylisporites mirabilis*
- Affinities (group): stem
- Affinities: *Cyathea*
- Order (of clade calibrated): Cyatheales
- Family (of clade calibrated): Cyatheaceae
- Notes on affinities: Based on the spore morphology (Plate V, fig. 10), with three equatorial pores that are characteristic of some *Cyathea* (i.e., *Cnemidaria*-type spores); directly compared to *Hemitelia karsteniana*. There are some doubts on the "holotype" referred in the original (see compilers preface of Traverse and Ames, 1979). In this case, the specimen number is the same for *Dicksonia dubia* and *Hemitelia mirabilis*. Possibly, this reflects the presence of multiple spores in the same sample.
- Reference: Bolchotinova (1953)

#### Fossil Locality

- Country: Russia
- Type locality: Southern Urals, left bank of the Ayat settlement, against the settlement of Novonikolayevskogo

#### Fossil Specimen

- Organs: Spores
- Specimen (Holotype): ИГН АН СССР No. 3527/41
- Collection: Mining-Geolical Institute of Western Siberia, Academy os Sciences, USSR (?)

---

### *Lindsaeaceae gen. et. sp. indet.*†

#### Calibration

- NFos: 73
- Minimum age: 93.9
- Node calibrated: stem Lindsaeaceae

#### Fossil Age

- Lower limit of oldest stratigraphic age: 100.5
- Upper limit of oldest stratigraphic age: 93.9
- Stratigraphic age: Upper Cretaceous (Cenomonian)
- Absolute age: 98.79 + 0.62 (U-Pb) (Cenomanian)
- Reference time scale: ICS (v2017/02)
- Notes on age: Burmese amber (Burmite), Earliest Upper Cretaceous, earliest Cenomanian, absolute age 98.79 ± 0.62 million years ago established by U-Pb dating of zircons from the rind of the unprocessed amber.

#### Fossil Identity

- Author: Regalado, A.R.Schmidt, Müller, Kobbert, H.Schneid. et Heinrichs
- Full taxon name: Lindsaeaceae gen. et. sp. indet. Regalado, A.R.Schmidt, Müller, Kobbert, H.Schneid. et Heinrichs
- Affinities (group): stem
- Affinities: Lindsaeaceae
- Order (of clade calibrated): Polypodiales
- Family (of clade calibrated): Lindsaeaceae
- Reference: Regalado et al. (2017)

#### Fossil Locality

- Country: Myanmar
- Type locality: Amber mines near Tanai, Ledo Road, 105 km northwest of Myitkyina, Kachin State, Myanmar
- Formation: Burmese amber (Burmite)

#### Fossil Specimen

- Organs: Fertile foliage, sporangia, spores
- Specimen (Holotype): GZG.BST.21952
- Collection: Geoscientific Collections of the Georg August University Go€ttingen

---

### *Lophosoria cupulatus*†

#### Calibration

- NFos: 74
- Minimum age: 113
- Node calibrated: stem *Lophosoria*

#### Fossil Age

- Lower limit of oldest stratigraphic age: 125
- Upper limit of oldest stratigraphic age: 113
- Stratigraphic age: Aptian
- Reference time scale: ICS (v2017/02)

#### Fossil Identity

- Author: Cantrill
- Full taxon name: *Lophosoria cupulatus* Cantrill
- Affinities (group): stem
- Affinities: *Lophosoria*
- Order (of clade calibrated): Cyatheales
- Family (of clade calibrated): Dicksoniaceae
- Notes on affinities: Affinities with the Lophosoriaceae, based on the distinctive spores of *Cyatheacidites annulatus*. The spores from *Lophosoria cupulatus* are best placed within *C. annulatus* Cookson ex Potonie, and confirm placement within the Lophosoriaceae.
- Reference: Cantrill (1998)

#### Fossil Locality

- Country: Antartica
- Type locality: Snow Island, Shetland Islands
- Formation: Cerro Negro Fm.

#### Fossil Specimen

- Organs: Fertile pinnule, sporangia, spores
- Specimen (Holotype): P.2501.69a (holotype); Plate V, figure 10
- Collection: British Antarctic Survey collections

---

### *Loxsomopteris anasilla*†

#### Calibration

- NFos: 79
- Minimum age: 125

- Node calibrated: stem Loxsomataceae

#### Fossil Age

- Lower limit of oldest stratigraphic age: 129.4
- Upper limit of oldest stratigraphic age: 125
- Stratigraphic age: Barremain or Aptian
- Reference time scale: ICS (v2017/02)
- Notes on age: Preliminary examination of pollen preparation of the fossil-bearing bed is consistent with its placement in Zone I, which is probably Aptian in age.

#### Fossil Identity

- Author: J.Skog
- Full taxon name: *Loxsomopteris anasilla* J.Skog
- Affinities (group): stem
- Affinities: Loxsomataceae
- Order (of clade calibrated): Cyatheales
- Family (of clade calibrated): Loxsomataceae
- Notes on affinities: On the basis of scletoric outer cortex and the characteristic hairs on the epidermis, the fossil is described as a new genus in the Losxomataceae.
- Reference: Skog (1976)

#### Fossil Locality

- Country: North America
- Type locality: Paint Branch, USNM locality 14212
- Formation: Potomac Group, Patuxent Fm.

#### Fossil Specimen

- Organs: Stem
- Specimen (Holotype): USNM 208539, a , b, c 1-21
- Collection: Paleobotany Collections, US National Museum of Natural History

---

### *Lygodium sanshuiense*†

#### Calibration

- NFos: 30
- Minimum age: 59.2
- Node calibrated: crown Lygodiaceae

#### Fossil Age

- Lower limit of oldest stratigraphic age: 61.6

- Upper limit of oldest stratigraphic age: 59.2
- Stratigraphic age: middle Paleocene (Selandian)
- Reference time scale: ICS (v2017/02)

**Fossil Identity**

- Author: Naugolnykh, Tu, Liu et Jin
- Full taxon name: *Lygodium sanshuiense* Naugolnykh, Tu, Liu et Jin
- Affinities (group): crown
- Affinities: Lygodiaceae
- Order (of clade calibrated): Schizaeales
- Family (of clade calibrated): Lygodiaceae
- Notes on affinities: Based upon their vegetative morphology, epidermal anatomy and the structure of the conducting tissues.
- Reference: Naugolnykh et al. (2020)

**Fossil Locality**

- Country: China
- Type locality: Jinfu Road, Sanshui District, Foshan City, Guangdong, China
- Formation: Buxin Fm

**Fossil Specimen**

- Organs: Sterile foliage
- Specimen (Holotype): SSBX-020
- Collection: Museum of Biology, Sun Yet-sen University, Guangzhou, China

---

### *Marattia aganzhenensis*†

**Calibration**

- NFos: 31
- Minimum age: 174.1
- Node calibrated: crown Marattiaceae

**Fossil Age**

- Lower limit of oldest stratigraphic age: 201.3
- Upper limit of oldest stratigraphic age: 174.1
- Stratigraphic age: Early Jurassic
- Reference time scale: ICS (v2017/02)

**Fossil Identity**

- Author: Yang, Wang et Pfefferkorn

- Full taxon name: *Marattia aganzhenensis* Yang, Wang et Pfefferkorn
- Affinities (group): crown
- Affinities: Marattiaceae
- Order (of clade calibrated): Marattiales
- Family (of clade calibrated): Marattiaceae
- Notes on affinities: The affinities are confirmed by the phylogenetic analysis of Rothwell et al. (2018), placing it within crown Marattiaceae.
- Reference: Yang et al. (2008)

**Fossil Locality**

- Country: China
- Type locality: Aganzhen, Lanzhou, Gansu, China
- Formation: Daxigou Fm.

**Fossil Specimen**

- Organs: Fertile foliage, sporangia, spores
- Specimen (Holotype): Holotype: NWGP-90001
- Collection: College of Earth and Environmental Sciences, Lanzhou University

---

### *Matonia jeffersonii*†

**Calibration**

- NFos: 116
- Minimum age: 100.5
- Node calibrated: stem *Matonia*

**Fossil Age**

- Lower limit of oldest stratigraphic age: 113
- Upper limit of oldest stratigraphic age: 100.5
- Stratigraphic age: late Albian
- Reference time scale: ICS (v2017/02)
- Notes on age: These fluvial strata have been assigned a Late Albian age on the basis of bracketing molluscan faunas (105 Ma).

**Fossil Identity**

- Author: Nagalingum et Cantrill
- Full taxon name: *Matonia jeffersonii* Nagalingum et Cantrill
- Affinities (group): stem
- Affinities: *Matonia*
- Order (of clade calibrated): Gleicheniales
- Family (of clade calibrated): Matoniaceae

- Notes on affinities: Assigned to Matoniaceae based on the presence of circular to oval sori with a peltate indusium.
- Reference: Nagalingum and Cantrill (2006)

**Fossil Locality**

- Country: Antarctica
- Type locality: British Antarctic Survey locality KG. 2816
- Formation: Triton Point Formation, Fossil Bluff Group

**Fossil Specimen**

- Organs: Sterile and fertile fronds, sporangia, spores
- Specimen (Holotype): KG. 2816.27b
- Collection: British Antarctic Survey

---

### *Neolepisorus chingii*†

**Calibration**

- NFos: 147
- Minimum age: 7.25
- Node calibrated: crown *Lepisorus*

**Fossil Age**

- Lower limit of oldest stratigraphic age: 11.63
- Upper limit of oldest stratigraphic age: 7.25
- Stratigraphic age: Late Miocene (Messinian-Tortonian)
- Reference time scale: ICS (v2017/02)
- Notes on age: Based on biostratigraphic and lithostratigraphic correlations.

**Fossil Identity**

- Author: Xie, Li, zhang, Shao, Wu et Sun
- Full taxon name: Neolepisorus chingii Xie, Li, zhang, Shao, Wu et Sun
- Affinities (group): crown
- Affinities: *Lepisorus*
- Order (of clade calibrated): Polypodiales
- Family (of clade calibrated): Polypodiaceae
- Notes on affinities: Characters including simple frond with a clear midrib, several prominent lateral veins somewhat zigzagging without reaching the margin, and simple or forked free-ending veinlets indicate affinities with *Lepisorus* sect. *Neolepisorus*. However, the fossil differs from extant taxa by number of sori (one line of sori on either side of the midrib in the fossil vs. many in extant taxa) and sorus shape (elliptic-fusiform in the fossil vs. orbicular in extant taxa), and *Neolepisorus* is nested within *Lepisorus* (as sect. *Neolepisorus* sensu Zhao

et al., 2020). Therefore we treat this conservatively as crown *Lepisorus* instead of assignment to sect. *Neolepisorus*.
- Reference: Xie et al. (2016)

**Fossil Locality**

- Country: China
- Type locality: Bangmai Basin in Lincang City of Yunnan Province
- Formation: Bangmai fm.

**Fossil Specimen**

- Organs: Fertile foliage
- Specimen (Holotype): MCD090225-037
- Collection: School of Earth Sciences, Lanzhou University

---

### *Onoclea sensibilis*

**Calibration**

- NFos: 93
- Minimum age: 56
- Node calibrated: stem *Onoclea*

**Fossil Age**

- Lower limit of oldest stratigraphic age: 59.2
- Upper limit of oldest stratigraphic age: 56
- Stratigraphic age: late Paleocene (Thanetian)
- Reference time scale: ICS (v2017/02)

**Fossil Identity**

- Author: L.
- Full taxon name: *Onoclea sensibilis* L.
- Affinities (group): stem
- Affinities: *Onoclea*
- Order (of clade calibrated): Polypodiales
- Family (of clade calibrated): Onocleaceae
- Reference: Rothwell and Stockey (1991)

**Fossil Locality**

- Country: North America
- Type locality: Munce's Hill, near Red Deer, Alberta
- Formation: Paskapoo Fm

#### Fossil Specimen

- Organs: Sterile and fertile fronds, sporangia, spores
- Specimen (Holotype): S17971-S18236, S18238-S19626, S23405-S23523, S23526-S23556, S23968-S23997, S26372-S26273, S26292-S26317, S26319-S26328, S26479-S26480
- Collection: University of Alberta Paleobotanical Collections (U.A.P.C.)

---

### *Ophioglossum senomanicum*†

#### Calibration

- NFos: 94
- Minimum age: 93.9
- Node calibrated: stem *Ophioglossum*

#### Fossil Age

- Lower limit of oldest stratigraphic age: 100.5
- Upper limit of oldest stratigraphic age: 93.9
- Stratigraphic age: Cenomanian-Turonian
- Reference time scale: ICS (v2017/02)

#### Fossil Identity

- Author: Chlonova
- Full taxon name: *Ophioglossum senomanicum* Chlonova
- Affinities (group): stem
- Affinities: *Ophioglossum*
- Order (of clade calibrated): Ophioglossales
- Family (of clade calibrated): Ophioglossaceae
- Notes on affinities: The spore bears a resemblance to modern *Ophioglossum lusitanicum* and *Ophioglossum falcatum*, differing from them in larger sizes and in the presence of a smooth area around the three-beam slit.
- Reference: Chlonova (1960)

#### Fossil Locality

- Country: Russia
- Type locality: Chulym, near the village of Suchkovo and 500 m from the village of Simonovo, right bank of the river Kemr, 2 km below the mouth
- Formation: Not provided

#### Fossil Specimen

- Organs: Spores
- Specimen (Holotype): Табл. III, фиг. 13—16

- Collection: Geological Institute of the Academy of Sciences of the USSR

---

### *Osmunda granulata*†

#### Calibration

- NFos: 95
- Minimum age: 93.9
- Node calibrated: stem *Osmunda*

#### Fossil Age

- Lower limit of oldest stratigraphic age: 100.5
- Upper limit of oldest stratigraphic age: 93.9
- Stratigraphic age: Cenomanian-Turonian
- Reference time scale: ICS (v2017/02)

#### Fossil Identity

- Author: (Maljavkina) Chlonova
- Full taxon name: *Osmunda granulata* (Maljavkina) Chlonova
- Affinities (group): stem
- Affinities: *Osmunda*
- Order (of clade calibrated): Osmundales
- Family (of clade calibrated): Osmundaceae
- Notes on affinities: The surface of the exine, covered with small papillae, makes it possible to compare this spore with the spores of modern ferns *Osmunda regalis* and *Osmunda claytoniana* (collection of reference spore preparations) and with *Osmunda triassica*. But none of these species can be identified as the same as the fossil.
- Reference: Chlonova (1960)

#### Fossil Locality

- Country: Russia
- Type locality: Chulym, near the village of Suchkovo and 500 m from the village of Simonovo, right bank of the river Kemr, 2 km below the mouth
- Formation: Not provided

#### Fossil Specimen

- Organs: Spores
- Specimen (Holotype): Табл. III, фиг. 4—5
- Collection: Geological Institute of the Academy of Sciences of the USSR

---

### *Paralygodium meckertii*†

#### Calibration

- NFos: 113
- Minimum age: 86.3
- Node calibrated: crown Schizaeaceae

#### Fossil Age

- Lower limit of oldest stratigraphic age: 89.8
- Upper limit of oldest stratigraphic age: 86.3
- Stratigraphic age: Coniacian
- Reference time scale: ICS (v2017/02)
- Notes on age: The Eden Main localities have been dated as upper Turonian to lower Coniacian using mollusc index fossils.

#### Fossil Identity

- Author: Karafit et Stockey
- Full taxon name: *Paralygodium meckertii* Karafit et Stockey
- Affinities (group): crown
- Affinities: Schizaeaceae
- Order (of clade calibrated): Schizaeales
- Family (of clade calibrated): Schizaeaceae
- Notes on affinities: Affinities with Schizaceae sensu lato, which includes *Lygodium*, *Anemia*, and *Schizea*; but directly compared to *Anemia*, thus assignment to Schizeaceae s.s.
- Reference: Karafit and Stockey (2008)

#### Fossil Locality

- Country: Canada
- Type locality: Eden Quarry
- Formation: Dunsmuir Member, Comox Formation

#### Fossil Specimen

- Organs: Fertile foliage, sporangia, spores
- Specimen (Holotype): P14457 E top
- Collection: University of Alberta Paleobotanical Collections (UAPC-ALTA)

---

### *Pleopeltis dominicensis*†

#### Calibration

- NFos: 33

- Minimum age: 15.97
- Node calibrated: crown *Pleopeltis*

#### Fossil Age

- Lower limit of oldest stratigraphic age: 20.44
- Upper limit of oldest stratigraphic age: 15.97
- Stratigraphic age: Burdigalian
- Reference time scale: ICS (v2017/02)
- Notes on age: The age of the Dominican amber deposit has been recently constrained from the stratigraphic age estimation of Burdigalian (15.8–20.3 Mya) to approximately 16 Ma.

#### Fossil Identity

- Author: H.Schneid., Heinrichs et A.R.Schmidt
- Full taxon name: *Pleopeltis dominicensis* H.Schneid., Heinrichs et A.R.Schmidt
- Affinities (group): crown
- Affinities: *Pleopeltis*
- Order (of clade calibrated): Polypodiales
- Family (of clade calibrated): Polypodiaceae
- Notes on affinities: The relationships of the fossil were explored by: (1) comparing morphological similarities with extant taxa; (2) identifying apomorphic character states by reconstructing the evolution of characters preserved in the fossil; and (3) analyses employing phylomorphospace reconstruction.
- Reference: Schneider et al. (2015)

#### Fossil Locality

- Country: Dominincan Republic
- Type locality: Burdigalian amber bearing strata of Dominican Republic
- Formation: Dominican amber deposit

#### Fossil Specimen

- Organs: Fertile foliage
- Specimen (Holotype): AMNH-DR-ASHS-1
- Collection: Dominican amber collection of the American Museum of Natural History, New York

---

### *Polypodium radonii*†

#### Calibration

- NFos: 97
- Minimum age: 27.82
- Node calibrated: stem *Polypodium* s.l.

#### Fossil Age

- Lower limit of oldest stratigraphic age: 33.9
- Upper limit of oldest stratigraphic age: 27.82
- Stratigraphic age: Early Oligocene
- Absolute age: 29-30 Mya (Rupelian)
- Reference time scale: ICS (v2017/02)
- Notes on age: Radiometric dating of the sodalite trachyte which penetrated into the sediments results in an age of 20-30 Mya for the fossiliferous shale.

#### Fossil Identity

- Author: Kvacek
- Full taxon name: *Polypodium radonii* Kvacek
- Affinities (group): stem
- Affinities: *Polypodium* s.l.
- Order (of clade calibrated): Polypodiales
- Family (of clade calibrated): Polypodiaceae
- Notes on affinities: The fossils are compared to *Polypodium*, with reference to *P. vulgare* complex. Our tree rendered a paraphyletic *Polypodium* by monotypic *Pleurosoriopsis*. This left us with two options: assign the fossil to *Polypodium* s.s. (excluding *Pleurosoriopsis* and *P. pellucidum*) or to *Polypodium* s.l. (including *Pleurosoriopsis* and *P. pellucidum*). As the fossil clearly aligns with *Polypodium*, we here chose to assign it to crown *Polypodium* s.l.
- Reference: Kvacek (2001)

#### Fossil Locality

- Country: Europe
- Type locality: Holy Kluk hill near ProboStov
- Formation: Usti Formation of the Ceske stredohori Mountains

#### Fossil Specimen

- Organs: Fertile foliage, sporangia, spores
- Specimen (Holotype): NM G 7765a
- Collection: National Museum, Praha

---

### *Polystichum pacltovae*†

#### Calibration

- NFos: 114
- Minimum age: 23.03
- Node calibrated: stem *Polystichum*

#### Fossil Age

- Lower limit of oldest stratigraphic age: 33.9
- Upper limit of oldest stratigraphic age: 23.03
- Stratigraphic age: Oligocene
- Reference time scale: ICS (v2017/02)

#### Fossil Identity

- Author: Kvacek
- Full taxon name: *Polystichum pacltovae* Kvacek
- Affinities (group): stem
- Affinities: *Polystichum*
- Order (of clade calibrated): Polypodiales
- Family (of clade calibrated): Dryopteridaceae
- Reference: Kvacek and Teodoridis (2020)

#### Fossil Locality

- Country: Czech Republic
- Type locality: Ústí nad Labem–Mojžíř
- Formation: Děčín Formation of the České středohoří Mts.

#### Fossil Specimen

- Organs: Sterile and fertile foliage, sporangia
- Specimen (Holotype): NM G 12419, NM G 12420
- Collection: Paleontological department of the National Museum in Prague

---

### *Protocalamostachys farringtonii*†

#### Calibration

- NFos: 64
- Minimum age: 346.7
- Node calibrated: stem Equisetales

#### Fossil Age

- Lower limit of oldest stratigraphic age: 358.9
- Upper limit of oldest stratigraphic age: 346.7
- Stratigraphic age: Late Tournaisian
- Reference time scale: ICS (v2017/02)
- Notes on age: The Oxroad Bay sequence has been dated as late Tournaisian. However, based on Scott and Galtier (1996), the Tournaisian age of these localities might be revised to Visean or even Late Visean.

**Fossil Identity**

- Author: Bateman
- Full taxon name: *Protocalamostachys farringtonii* Bateman
- Affinities (group): stem
- Affinities: Equisetales
- Order (of clade calibrated): Equisetales
- Notes on affinities: Phylogenetic analyses of Elgorriaga et al. (2018) place this fossil as sister to extant Equisetaceae.
- Reference: Bateman (1991)

**Fossil Locality**

- Country: UK
- Type locality: Oxroad Bay, East Lothian (horizon 2)
- Formation: Dinantian of Scottish Midland Valley

**Fossil Specimen**

- Organs: Strobilus, sporangiophores, sporagia, spores
- Specimen (Holotype): Syntypes: OBC001d1B, OBC061b1T, OBC084dB, OBC084eT, OBC084gB
- Collection: British Museum (Natural History)

---

### *Protodrynaria takhtajanii*†

**Calibration**

- NFos: 125
- Minimum age: 33.9
- Node calibrated: stem *Drynaria*

**Fossil Age**

- Lower limit of oldest stratigraphic age: 37.8
- Upper limit of oldest stratigraphic age: 33.9
- Stratigraphic age: Eocene-Oligocene (Priabonian)
- Reference time scale: ICS (v2017/02)

**Fossil Identity**

- Author: Vikulin et A.Bobr.
- Full taxon name: *Protodrynaria takhtajanii* Vikulin et A.Bobr.
- Affinities (group): stem
- Affinities: *Drynaria*
- Order (of clade calibrated): Polypodiales

- Family (of clade calibrated): Polypodiaceae
- Notes on affinities: The assignment to *Drynaria* is confirmed by the phylogenetic analyses of Su et al. (2011); ultimately Su et al. (2011) state that ‘our results are in favor of drynarioid affinities for *P. takhtajanii*’, thus we assign the fossil as stem.
- Reference: Vikulin (1982)

#### Fossil Locality

- Country: Russia
- Type locality: The village Tim, Kursk Province
- Formation: Not provided

#### Fossil Specimen

- Organs: Fertile foliage
- Specimen (Holotype): collection 6842 (= 2035), specimen 2
- Collection: Chernyshev Central Scientific Geological and Prospecting Museum

---

### *Radstockia kidsonii*†

#### Calibration

- NFos: 86
- Minimum age: 307
- Node calibrated: stem Marattiaceae

#### Fossil Age

- Lower limit of oldest stratigraphic age: 315.2
- Upper limit of oldest stratigraphic age: 307
- Stratigraphic age: Middle Pennsylvanian
- Reference time scale: ICS (v2017/02)

#### Fossil Identity

- Author: Taylor
- Full taxon name: *Radstockia kidsonii* Taylor
- Affinities (group): stem
- Affinities: Marattiaceae
- Order (of clade calibrated): Marattiales
- Family (of clade calibrated): Marattiaceae
- Notes on affinities: The affinities are confirmed by the phylogenetic analysis of Rothwell et al. (2018), placing it as stem Marattiaceae.
- Reference: Taylor (1967)

**Fossil Locality**

- Country: North America
- Type locality: Mazon Creek, Will Country, Illinois
- Formation: Francis Creek Shale, Carbondale Fm, Kewanee group

**Fossil Specimen**

- Organs: Sterile fronds, sporangia, spores
- Specimen (Holotype): Plate 6, fig. 1 (No. 1004)
- Collection: Peabody Museum of Natural History, Yale University, Paleobotanical collections

---

### *Regnellidium upatoensis*†

**Calibration**

- NFos: 98
- Minimum age: 83.6
- Node calibrated: stem *Regnellidium*

**Fossil Age**

- Lower limit of oldest stratigraphic age: 86.3
- Upper limit of oldest stratigraphic age: 83.6
- Stratigraphic age: Santonian
- Reference time scale: ICS (v2017/02)
- Notes on age: Biostratigraphic correlation places these outcrops in the Eutaw Formation or its non-marine equivalent on the Gulf Coastal Plain.

**Fossil Identity**

- Author: Lupia, Schneider, et Moeser
- Full taxon name: *Regnellidium upatoensis* Lupia, Schneider, et Moeser
- Affinities (group): stem
- Affinities: *Regnellidium*
- Order (of clade calibrated): Salviniales
- Family (of clade calibrated): Marsileaceae
- Reference: Lupia et al. (2000)

**Fossil Locality**

- Country: North America
- Type locality: Upatoi Creek, Chattahoochee County, Georgia, U.S.A
- Formation: Eutaw Formation

#### Fossil Specimen

- Organs: Sporocarp, megaspores
- Specimen (Holotype): PP45948
- Collection: paleobotanical collections of the Field Museum, Chicago

---

### *Regnellites nagashimae*†

#### Calibration

- NFos: 87
- Minimum age: 145
- Node calibrated: stem Marsileaceae

#### Fossil Age

- Lower limit of oldest stratigraphic age: 163.5
- Upper limit of oldest stratigraphic age: 145
- Stratigraphic age: Upper Jurassic to Berrasian
- Reference time scale: ICS (v2017/02)

#### Fossil Identity

- Author: Yamada et Kato
- Full taxon name: *Regnellites nagashimae* Yamada et Kato
- Affinities (group): stem
- Affinities: Marsileaceae
- Order (of clade calibrated): Salviniales
- Family (of clade calibrated): Marsileaceae
- Notes on affinities: Phylogenetic analyses place it as a stem relative, sister to extant Marsileaceae; although there is low support (BP = 68) for this position, rendering the possibility of the fossil representing a crown member of Salviniales.
- Reference: Yamada and Kato (2002)

#### Fossil Locality

- Country: Japan
- Type locality: Locality 102 on Nakao Forestry Road, Shimonoseki, Yamaguchi Prefecture
- Formation: Kiyosue Fm

#### Fossil Specimen

- Organs: Rhizomes, petioles, sporocarps, spores
- Specimen (Holotype): Holotype. NSM-PP-9944
- Collection: National Science Museum Tokyo

---

### *Stachypteris spicans*†

#### Calibration

- NFos: 80
- Minimum age: 168.3
- Node calibrated: stem *Lygodium*

#### Fossil Age

- Lower limit of oldest stratigraphic age: 170.3
- Upper limit of oldest stratigraphic age: 168.3
- Stratigraphic age: Bajocian
- Reference time scale: ICS (v2017/02)

#### Fossil Identity

- Author: Pomel
- Full taxon name: *Stachypteris spicans* Pomel
- Affinities (group): stem
- Affinities: *Lygodium*
- Order (of clade calibrated): Schizaeales
- Family (of clade calibrated): Lygodiaceae
- Notes on affinities: Comparing the Jurassic *Klukia exilis* and *Stachypteris spicans* spores with Recent spores, one may say that they are similar but not strictly comparable, especially their greatly reduced proximal sculpture that has not been seen in the Recent spores. Within the Schizaeaceae, however, they certainly resemble most closely reticulate *Lygodium* spores (van Konijnenburg-van Cittert, 1989); therefore these may mark the split between *Lygodium* and the *Anemia*/*Schizaea* clade.
- Reference: Harris (1961)

#### Fossil Locality

- Country: UK
- Type locality: Marske Quarry, Hasty Bank and Whitby, Yorkshire
- Formation: Lower Deltaic Series, Yorkshire flora

#### Fossil Specimen

- Organs: Sterile and fertile fronds, sporangia, spores
- Specimen (Holotype): V.27708a-b, V.31908a, V.27709a, V.31910a
- Collection: British Museum (Natural History)

### *Szea sinensis*†

#### Calibration

- NFos: 68
- Minimum age: 272.95
- Node calibrated: stem Gleicheniaceae

#### Fossil Age

- Lower limit of oldest stratigraphic age: 283.5
- Upper limit of oldest stratigraphic age: 272.95
- Stratigraphic age: late Early Permian (Kungurian-Roadian)
- Reference time scale: ICS (v2017/02)
- Notes on age: Equivalent to Guadalupian sediments of North America.

#### Fossil Identity

- Author: Yao et Taylor
- Full taxon name: *Szea sinensis* Yao et Taylor
- Affinities (group): stem
- Affinities: Gleicheniaceae
- Order (of clade calibrated): Gleicheniales
- Family (of clade calibrated): Gleicheniaceae
- Reference: Zhaoqi and Taylor (1988)

#### Fossil Locality

- Country: China
- Type locality: Funiushan Coal Mine, between Nan- jing and Zhenjiang, approximately 20 km from Zhenjiang, Jiangsu Province
- Formation: Lower portion of Lungtan Formation.

#### Fossil Specimen

- Organs: Fertile foliage, sporangia, spores
- Specimen (Holotype): Specimen PB 9270
- Collection: Palaeobotany Collection, Nanjing Institute of Geology and Palaeontology, Academia Sinica, Nanjing, Jiangsu Province, China

---

### *Thelypteris goldianum*†

#### Calibration

- NFos: 129
- Minimum age: 72.1

- Node calibrated: crown Thelypteridaceae

**Fossil Age**

- Lower limit of oldest stratigraphic age: 83.6
- Upper limit of oldest stratigraphic age: 72.1
- Stratigraphic age: Campanian
- Reference time scale: ICS (v2017/02)

**Fossil Identity**

- Author: (Lesquereux) Crabtree
- Full taxon name: *Thelypteris goldianum* (Lesquereux) Crabtree
- Affinities (group): crown
- Affinities: Thelypteridaceae
- Order (of clade calibrated): Polypodiales
- Family (of clade calibrated): Thelypteridaceae
- Reference: Crabtree (1987)

**Fossil Locality**

- Country: United States
- Type locality: A horizon at site 42
- Formation: Two Medicine Formation

**Fossil Specimen**

- Organs: Fertile foliage, sporangia
- Specimen (Holotype): 2A/F5/001
- Collection: University of Montana Paleontological Museum

---

### *Tomaniopteris katonii*†

**Calibration**

- NFos: 90
- Minimum age: 242
- Node calibrated: stem Matoniaceae

**Fossil Age**

- Lower limit of oldest stratigraphic age: 247.2
- Upper limit of oldest stratigraphic age: 242
- Stratigraphic age: early Middle Triassic (Anisian)
- Reference time scale: ICS (v2017/02)

- Notes on age: The peat occurs in carbonaceous mudstones of the upper Fremouw Formation, which is early Middle Triassic in age based on palynology and vertebrates.

**Fossil Identity**

- Author: Klavins, Taylor et Taylor
- Full taxon name: *Tomaniopteris katonii* Klavins, Taylor et Taylor
- Affinities (group): stem
- Affinities: Matoniaceae
- Order (of clade calibrated): Gleicheniales
- Family (of clade calibrated): Matoniaceae
- Notes on affinities: Combination of characters that affiliates them with the Matoniaceae.
- Reference: Klavins et al. (2004)

**Fossil Locality**

- Country: Antarctica
- Type locality: Fremouw Peak, Queen Alexandra Range, central Transantarctic Mountains
- Formation: Upper Fremouw Formation

**Fossil Specimen**

- Organs: Sori, sporangia
- Specimen (Holotype): Holotype, 49 slides of specimen 11248A, slide Nos. 20461–20465, 20466–20510
- Collection: Division of Paleobotany of the Natural History Museum and Biodiversity Research Center, University of Kansas

---

### *Woodwardia sp. indet.*†

**Calibration**

- NFos: 123
- Minimum age: 72.1
- Node calibrated: stem *Woodwardia*

**Fossil Age**

- Lower limit of oldest stratigraphic age: 83.6
- Upper limit of oldest stratigraphic age: 72.1
- Stratigraphic age: Campanian
- Absolute age: 76.1–72.5 Ma (Amato_etal-2017)
- Reference time scale: ICS (v2017/02)
- Notes on age: Tentative age of late (but not latest) Maastrichtian of the Jose Creeek Member based on stratigraphic correlations.

**Fossil Identity**

- Author: Upchurch et Mack
- Full taxon name: *Woodwardia sp.* indet. Upchurch et Mack
- Affinities (group): stem
- Affinities: *Woodwardia*
- Order (of clade calibrated): Polypodiales
- Family (of clade calibrated): Blechnaceae
- Notes on affinities: cf. the fossils from Crabtree (1987)
- Reference: Upchurch Jr. and Mack (1998)

**Fossil Locality**

- Country: United States
- Type locality: Not provided
- Formation: Jose Creek Member of the McRae Formation

**Fossil Specimen**

- Organs: Sterile frond
- Specimen (Holotype): Figure 1
- Collection: Not provided

---
